# The Autophagy Receptor TAX1BP1 (T6BP) is a novel player in antigen presentation by MHC-II molecules

**DOI:** 10.1101/2021.04.21.440798

**Authors:** Mathias Pereira, Clémence Richetta, Gabriela Sarango, Anita Kumari, Michael Ghosh, Lisa Bertrand, Cédric Pionneau, Morgane Le Gall, Sylvie Grégoire, Raphaël Jeger-Madiot, Elina Rosoy, Frédéric Subra, Olivier Delelis, Mathias Faure, Audrey Esclatine, Stéphanie Graff-Dubois, Stefan Stevanović, Bénédicte Manoury, Bertha Cecilia Ramirez, Arnaud Moris

**Affiliations:** Université Paris-Saclay, CEA, CNRS, Institute for Integrative Biology of the Cell (I2BC), 91190, Gif-sur-Yvette, France; Sorbonne Université, INSERM, CNRS, Center for Immunology and Microbial Infections (CIMI-Paris), 75013, Paris, France; Department of Immunology, Institute for Cell Biology, University of Tübingen, 72076, Tübingen, Germany; Sorbonne Université, INSERM, UMS Production et Analyse de données en Sciences de la vie et en Santé, PASS, Plateforme Post-génomique de la Pitié Salpêtrière, 75013, Paris, France; 3P5 proteom’IC facility, Université de Paris, Institut Cochin, INSERM U1016, CNRS-UMR 8104, 75014, Paris, France; LBPA, ENS-Paris Saclay, CNRS UMR8113, Université Paris Saclay, Gif-sur-Yvette, France; CIRI, Centre International de Recherche en Infectiologie, Université de Lyon, Inserm U1111, Université Claude Bernard Lyon 1, CNRS, UMR5308, ENS de Lyon, F-69007, Lyon, France; Equipe Labellisée par la Fondation pour la Recherche Médicale, FRM; Institut Necker Enfants Malades, INSERM U1151-CNRS UMR 8253, Faculté de médecine Necker, Université de Paris, 75015, Paris, France

**Author notes:** **To whom correspondence should be addressed:** Arnaud Moris, or Benedicte Manoury. These authors contributed equally to this work (Mathias Pereira, Clémence Richetta, Gabriela Sarango). SGD and RJM: Sorbonne Université, INSERM U959, Immunology-Immunopathology-Immunotherapy (i3), Paris, France.

**Keywords:** Calnexin, CD4^+^ T cell activation, Interactome, Immunopeptidome, Virus

## Abstract

CD4^+^ T lymphocytes play a major role in the establishment and maintenance of immunity. They are activated by antigenic peptides derived from extracellular or newly synthesized (endogenous) proteins presented on the surface of antigen presenting cells (APCs) by the MHC-II molecules. The pathways leading to endogenous MHC-II presentation remain poorly characterized. We demonstrate here that the autophagy receptor, T6BP, influences both autophagy-dependent and -independent endogenous presentation of HIV- and HCMV-derived peptides. By studying the immunopeptidome of MHC-II molecules, we show that T6BP affects both the quantity and quality of peptides presented. T6BP silencing induces the mislocalization of the MHC-II-loading compartments and a rapid degradation of the invariant chain (CD74) without altering the expression and internalization kinetics of MHC-II molecules. We determined the interactome of T6BP, identified calnexin as a T6BP partner and show that CANX cytosolic tail is required for this interaction. Remarkably, calnexin silencing replicates the functional consequences of T6BP silencing: decreased CD4^+^ T cell activation and exacerbated CD74 degradation. Altogether, we unravel T6BP as a key player of the MHC-II-restricted endogenous presentation pathway and we propose one potential mechanism of action.

## Introduction

CD4^+^ helper T cells that orchestrate adaptive immune responses recognize pathogen- or tumour-derived peptides presented by the major histocompatibility complex class-II (MHC-II) molecules. MHC-II molecules are expressed by professional antigen-presenting cells (APC) such as B cells, macrophages and dendritic cells (DC), thymic epithelial cells (TEC), and by non-professional APCs in inflammatory conditions (Roche & Furuta, 2015; Wijdeven *et al*, 2018). The MHC-II transactivator, CIITA, governs the transcription of the MHC-II locus that includes genes encoding for the α- and β-chains of MHC-II molecules, the invariant chain Ii (CD74) and the chaperon proteins HLA-DM/HLA-DO (Reith *et al*, 2005). The transmembrane α- and β-chains are assembled within the endoplasmic reticulum (ER), where they associate with CD74 leading to the formation of nonameric αβ-CD74 complexes that traffic into late endo-lysosomal compartments named MIIC (Bakke & Dobberstein, 1990; Lotteau *et al*, 1990; Neefjes *et al*, 1990; Roche *et al*, 1991). In the MIIC, CD74 is progressively cleaved by vesicular proteases (Manoury *et al*, 2003; Nakagawa *et al*, 1998; Riese *et al*, 1996; Shi *et al*, 2000), leaving a residual MHC-II-associated Ii peptide (CLIP) that occupies the peptide binding groove (Bijlmakers *et al*, 1994; Busch *et al*, 1996; Roche & Cresswell, 1991). HLA-DM then facilitates the exchange of the CLIP fragments with high affinity peptides generated from pathogen- or tumor-derived antigens (Denzin & Cresswell, 1995; Morris *et al*, 1994; Sanderson *et al*, 1994). MHC-II molecules are then transported to the plasma membrane to expose antigenic peptides to CD4^+^ T cells (Thibodeau *et al*, 2019).

MHC-II molecules present peptides derived from extra- and intra-cellular sources of antigens, so-called exogenous and endogenous presentation, respectively (Veerappan Ganesan & Eisenlohr, 2017; Watts, 2004). Extracellular antigens are captured and internalized into APCs by various means including macropinocytosis, phagocytosis or receptor-mediated endocytosis (Roche & Furuta, 2015). Antigens are then delivered to the MIIC where they are progressively degraded by endo-lysosomal proteases such as cathepsins (Watts, 2004), into peptides (or epitopes) ranging from 12 to 25 amino acids in length, that can be loaded on nascent MHC-II molecules (Rudensky *et al*, 1991; Unanue *et al*, 2016). Epitopes from extracellular antigens can also bind, in early endosomes, to recycling MHC-II molecules (Pinet *et al*, 1995; Sinnathamby & Eisenlohr, 2003). The endogenous pathway relies on protein antigen synthesis by virus-infected (Eisenlohr & Hackett, 1989; Jacobson *et al*, 1988; Jaraquemada *et al*, 1990; Nuchtern *et al*, 1990; Sekaly *et al*, 1988; Thiele *et al*, 2015) or tumor cells (Tsuji *et al*, 2012). Some early *in vitro* studies showed that neosynthesized self-epitopes are displayed, after lysosomal proteolysis, by MHC-II molecules leading to CD4^+^ T cell activation (Bikoff & Birshtein, 1986; Rudensky & Yurin, 1989; Weiss & Bogen, 1989). More recently, it was shown that the initiation of CD4^+^ T cell responses to *influenza virus* is mainly driven by epitopes derived from the processing of intracellular antigens within APCs (Miller *et al*, 2015). However, the pathways leading to the loading of MHC-II molecules with endogenous antigens remain poorly characterized. Components of the MHC class-I (MHC-I) processing pathway such as proteasome have been implicated (Lich *et al*, 2000; Tewari *et al*, 2005). One unresolved issue is how cytosolic antigens are transported into MHC-II-enriched compartments (Crotzer & Blum, 2008; Dani *et al*, 2004). For some specific epitopes but not others, the transporter associated with antigen presentation of MHC-I molecules (TAP) has been linked with the delivery of endogenous peptide on MHC-II molecules (Malnati *et al*, 1992; Tewari *et al*., 2005). In fact, depending on the cellular localization, the trafficking and the nature of the antigen itself, different pathways might be involved in the degradation and delivery of endogenous antigens to MHC-II loading compartments (Leung, 2015; Mukherjee *et al*, 2001; Tewari *et al*., 2005).

The pathways of autophagy contribute to the processing of MHC-II-restricted endogenous antigens. The receptor of chaperone-mediated autophagy, LAMP-2A, has been shown to facilitate the presentation of a cytosolic self-antigen by MHC-II molecules (Zhou *et al*, 2005). The analysis of the MHC-II immunopeptidome revealed that macroautophagy (herein referred to as autophagy) also contributes to the processing of cytoplasmic and nuclear antigens (Dengjel *et al*, 2005). Autophagy is a self-eating cellular degradation pathway, in which double-membrane autophagosomes deliver their cytoplasmic constituents for lysosomal degradation (Kirkin, 2020). Using various models, several labs established that autophagy participates, in TECs, in the generation of MHC-II-restricted endogenous epitopes and strongly influences thymic selection of auto-reactive CD4^+^ T cells (Aichinger *et al*, 2013; Schuster *et al*, 2015). Other evidence that autophagy plays a role in endogenous antigen presentation comes from *in-vitro* studies using APCs transfected with mRNAs encoding tumor antigens (Dorfel *et al*, 2005) or cDNAs encoding tumor or viral antigens targeted to autophagosomes (Coulon *et al*, 2016; Fonteneau *et al*, 2016; Jin *et al*, 2014; Schmid *et al*, 2007). Targeting antigens to autophagosomes through the fusion to LC3, an autophagy effector that incorporates into and participates in the elongation of autophagosomes, enhances the capacity of APCs to activate antigen-specific CD4^+^ T cells (Coulon *et al*., 2016). However, overall, there are few examples where endogenous degradation of native tumor or viral antigens has been shown to be dependent on autophagy (Leung, 2015; Paludan *et al*, 2005). In fact, autophagy effectors may directly or indirectly affect the presentation of MHC-II-restricted antigens (Fletcher *et al*, 2018) by regulating, for instance, the delivery of proteases into the MIIC (Lee *et al*, 2010). Thereafter, autophagy also contributes to exogenous presentation of viral and bacterial antigens (Blanchet *et al*, 2010; Jagannath *et al*, 2009). The molecular links between autophagy and MHC-II-restricted antigen presentation, in particular the mechanisms allowing the delivery of autophagy-degraded antigens to the MIIC, are poorly defined.

A growing body of evidence indicates that autophagosomes selectively target their cargos, while excluding the rest of the cytoplasmic content (Kirkin, 2020). Several forms of selective autophagy exist, depending on the substrate, but all rely on the so-called autophagy receptors (ARs) that include: Nuclear Dot Protein 52 (NDP52), Optineurin (OPTN), Sequestosome-1 / p62, Next to BRCA1 gene protein-1 (NBR1) and TAX1-Binding Protein-1 also called TRAF6-Binding Protein (TAX1BP1/T6BP) (Kirkin & Rogov, 2019). ARs contain ubiquitin (Ub) and LC3-binding domains that allow on the one hand, binding to ubiquitinated proteins and on the other hand, their targeting into autophagosomes through interaction with LC3 on the internal membranes of forming autophagosomes (Kirkin & Rogov, 2019). As such, ARs are involved in multiple cellular processes including selective degradation of incoming bacteria and of damaged mitochondria, processes called xenophagy (Tumbarello *et al*, 2015) and mitophagy (Randow & Youle, 2014), respectively. In addition to their role in selective autophagy, T6BP, NDP52 and OPTN are required for the maturation of autophagosomes (Tumbarello *et al*, 2012). Thanks to the binding to myosin-VI, these ARs bridge autophagosomes to Tom-1-expressing endosomes and lysosomes, thus facilitating their fusion (Morriswood *et al*, 2007; Sahlender *et al*, 2005; Tumbarello *et al*., 2012). T6BP, OPTN, and NDP52, by promoting autophagosome maturation (Verlhac *et al*, 2015), are essential for the degradation of *Salmonella typhimurium* (Lin *et al*, 2019; Thurston *et al*, 2009; Wild *et al*, 2011). Remarkably, T6BP, NDP52, and p62 were also shown to orchestrate the maturation of early endosomes into late endosomes (Jongsma *et al*, 2016). This process also involves the Ub-binding domain of these receptors (Jongsma *et al*., 2016). Therefore, ARs exert multiple redundant but also exclusive roles in selective autophagy and in the traffic and maturation of vesicles such as autophagosomes and endosomes.

Here, we hypothesize that ARs may contribute at various, so far unknown, levels to MHC-II-restricted viral antigen presentation. We show that silencing of NDP52, OPTN and p62 in model APCs does not significantly affect the presentation of an autophagy-dependent antigen to CD4^+^ T cells. In contrast, T6BP influences both autophagy-dependent and -independent endogenous viral, as well as cellular, antigen processing and presentation by MHC-II molecules. In fact, the action of T6BP is not limited to viral antigens since the global repertoire of peptides presented by MHC-II molecules (immunopeptidome) is dramatically changed upon T6BP silencing. We show that T6BP silencing does not perturb the global cell-surface expression nor internalization kinetics of MHC-II molecules. However, it induces a significant relocalization of the MIIC closer to the nucleus and the generation of unstable MHC-II-peptide complexes. Importantly, we demonstrate that the absence of T6BP expression induces a strong and rapid degradation of the invariant chain CD74, which directly influences the quality of the peptide repertoire loaded on MHC-II molecules. Finally, to get a hint on possible mechanisms, we defined the interactome of T6BP and identified novel protein partners that potentially participate to the T6BP-mediated regulation of MHC-II peptide loading complex. Among them, we identified the ER chaperone calnexin (CANX). We show that T6BP binds the cytoplasmic tail of CANX known to regulate its ER functions. Finally, we provide the direct demonstration that silencing CANX also induces CD74 degradation and decreases the capacity of model APCs to activate CD4^+^ T cells. Altogether, this study unravels a new role for T6BP as a key player in MHC-II-restricted antigen presentation, and in CD4^+^ T cell immunity.

## Results

### T6BP silencing influences endogenous viral antigen presentation and CD4^+^ T cell activation

Owing to their functions in selective autophagy as well as in the maturation of autophagosomes, we focused our work on NDP52, OPTN, and T6BP asking whether these ARs might be involved in endogenous antigen presentation by MHC-II molecules and subsequent activation of CD4^+^ T cells. To this end, HeLa cells modified to express CIITA (HeLa-CIITA) were silenced for the expression of ARs using siRNAs targeting NDP52, OPTN and T6BP and evaluated for their capacity to activate CD4 T cell clones (Fig 1A). An siRNA targeting p62 was also included as this AR plays, in multiple models, a dominant role in selective autophagy but does not participate in the maturation of autophagosomes (Tumbarello *et al*., 2012). 24h post-siRNA transfection HeLa-CIITA cells were transfected with a plasmid encoding HIV-Gag protein fused to LC3. This Gag-LC3 fusion enables a specific targeting of Gag into autophagosomes and enhances HIV-specific T cell activation in an autophagy-dependent manner (Coulon *et al*., 2016). 48h post-siRNA treatment (24h post DNA transfection), we analysed by flow cytometry the percentages of living and of Gag-positive (Gag^+^) cells, using a viability dye and Gag intracellular staining, respectively (Fig S1A). In all tested conditions, the levels of Gag^+^ cells were similar and the sequential transfections (siRNA and cDNA) had no significant influence on cell viability (Fig S1B). The silencing of AR expression was also analysed by Western Blot. As compared to the control siRNA (CTRL), all siRNAs led to a marked decrease of AR expression (Fig 1C). HeLa-CIITA cells were then co-cultured with Gag-specific CD4^+^ T cells that we previously isolated and characterized (Moris *et al*, 2006). These Gag-specific CD4^+^ T cell clones recognize HIV-infected cells (Coulon *et al*., 2016; Moris *et al*., 2006). CD4+ T cell activation was monitored using IFN-γ-ELISPOT (Fig 1B). Cells transfected with CTRL, NDP52, OPTN, or p62 targeting siRNAs led to similar levels of CD4+ T cell activation (Fig 1B, right and left panel). In contrast, T6BP silencing greatly decreased the activation of Gag-specific T cells (Fig 1B, left panel). On average, T6BP silencing led to a 75% decrease of Gag-specific CD4^+^ T cell activation (Fig 1B, right panel).

**Figure 1.**
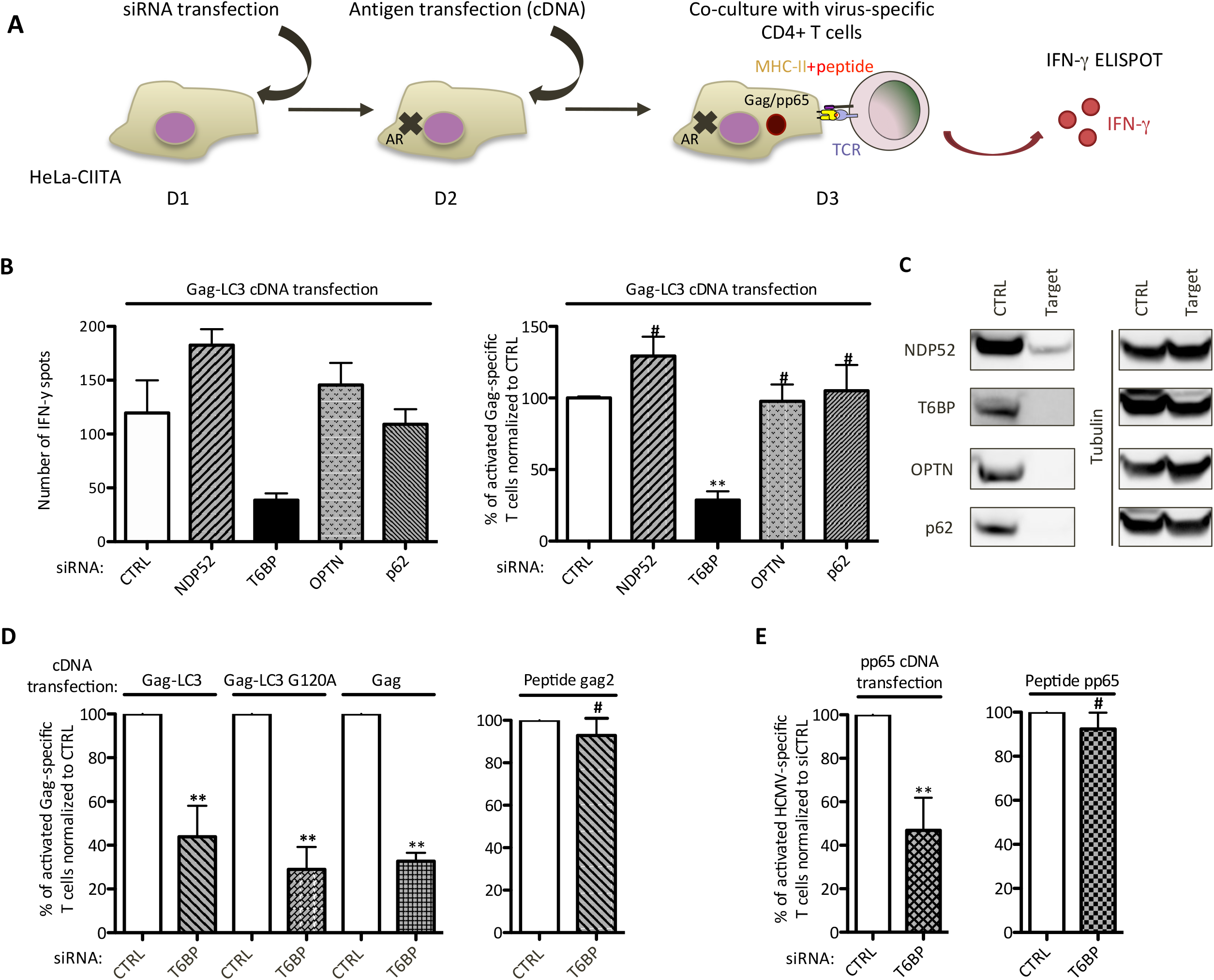
T6BP silencing decreases endogenous viral Ag presentation and CD4+ T cell activation. **(A)** Schematic representation of the experiment. Hela-CIITA cells were transfected with siRNAs targeting ARs and 24h later with plasmids encoding the antigens Gag, Gag-LC3, Gag-LC3 G120A or pp65. 24h post-DNA transfection, the HeLa-CIITA cells were co-cultured with antigen-specific CD4+ T cells and T cell activation was assessed using IFNγ-ELISPOT. AR: Autophagy Receptor; D: day; TCR: T-cell receptor. **(B)** Monitoring of Gag-specific T cell activation. HeLa-CIITA cells were treated as indicated above and transfected with a plasmid encoding Gag-LC3. Left panel, a representative experiment is shown. Right panel, three independent experiments are combined and presented as mean percentage (+/-SD). The y-axis represents the relative percentage of IFNγ spots reported to the secretion of IFNγ by the CD4+ T cell clones incubated with the siCTRL-treated HeLa-CIITA and set to 100%. The mean T cell activation levels of the three experiments using siCTRL or siT6BP were (100, 195, 119) and (35, 53, 38) IFNγ+ spots, respectively. 5000 T cell clones were seeded per well in technical triplicates. **(C)** 48h post-transfection of HeLa-CIITA cells with siRNAs targeting NDP52, T6BP, OPTN, and p62, AR expression was analyzed using Western Blot. Tubulin was used as control. The results are representative of at least 3 independent experiments and correspond to AR expression levels of the experiment in Figure 1B, left panel. **(D)** Left panel, as in Figure 1B right panel, but using DNA encoding Gag-LC3, Gag-LC3G120A, or Gag. With Gag-LC3, Gag-LC3 G120A or Gag antigens, the mean T cell activation levels using siCTRL or siT6BP ranged from (199-384), (91-141) and (343-415) or (98-273), (19-50) and (97-148) IFNγ+ spots, respectively. 5000 T cell clones were seeded per well in technical triplicates. Right panel, influence of T6BP silencing on peptide presentation by Hela-CIITA cells. The cognate peptide was added exogenously (gag2, 0,1µg/mL) on siRNA-treated cells (2h, 37°C), washed and T cell activation monitored using IFNγ-ELISPOT. The mean T cell activation levels using siCTRL or siT6BP were (100, 45, 42) and (71, 45, 33) IFNγ+ spots, respectively. 1000 T cell clones were seeded per well in technical triplicates. Three independent experiments are combined and presented as mean percentage (+/-SD) **(E)** As in (D) but using cDNA encoding HCMV pp65 antigen (left panel) or pp65 peptide (0,5µg/mL; right panel) and a pp65-specific CD4+ T cell line. Results of two independent experiments are represented. The mean T cell activation levels using siCTRL or siT6BP and pp65 DNA were (66, 106) and (39, 50) IFNγ+ spots, respectively. For the peptide: siCTRL or siT6BP: (185, 317) and (163, 292) IFNγ+ spots, respectively. 1000 T cell clones were seeded per well in technical triplicates. For all ELISPOT experiments, the background secretions of IFNγ by CD4+ T cells co-cultured with mock-treated HeLa-CIITA cells were used as negative controls and subtracted. CTRL: control. Wilcoxon’s tests; the symbols correspond to **:p<0.01; *:p<0.05 and #:p>0.05 comparing each experimental conditions solely with its internal control (siCTRL).

We next analysed the effect of T6BP silencing on autophagy-independent endogenous viral antigen processing by MHC-II molecules. As previously, HeLa-CIITA cells were first transfected with CTRL- or T6BP-targeting siRNAs (siCTRL and siT6BP respectively), then transfected with various plasmids encoding Gag, Gag-LC3_G120A_ or Gag-LC3, and co-cultured with the Gag-specific CD4^+^ T cells (Fig 1D). Importantly, in our previous work we demonstrated that newly synthetized Gag is processed in an autophagy independent manner both in monocyte-derived DCs (MDDCs) and in HeLa-CIITA cells (Coulon *et al*., 2016). This was shown by using productively infected MDDCs and Gag-cDNA-transfected HeLa-CIITA cells treated with various drugs influencing autophagy (e.g. 3-MA, Spautin-1 and Torin-1), shRNA targeting LC3 or overexpressing a trans-dominant mutant of Atg4B (Atg4BC74A), which blocks the formation of autophagosomes. Using the same tools, we also showed that the fusion with LC3 (Gag-LC3) targets Gag to autophagosome-mediated degradation leading to an enhancement of CD4^+^ T cell activation. Gag-LC3_G120A_ was also used as a negative control for autophagy-dependent degradation, as the G120A mutation in the C-terminus of LC3 abolishes the lipidation and incorporation of LC3 in the nascent membranes of autophagosomes, thus preventing Gag targeting into autophagosomes. Finally, we showed that upon Gag-LC3_G120A_-transfection, Gag antigens are processed in an autophagy independent manner (Coulon *et al*., 2016). 24h-post transfection and prior co-culture with the CD4^+^ T cells, the percentage of Gag^+^ cells and Gag expression levels and the cell viability were similar in all tested conditions (Fig S1C). As previously, T6BP silencing strongly decreased the capacity of HeLa-CIITA cells expressing Gag-LC3 to activate the Gag-specific T cells (Fig 1D, left panel). However, we observed that the effect of T6BP silencing was not limited to Gag-LC3 as the capacity of HeLa-CIITA expressing Gag- or Gag-LC3_G120A_ to activate the CD4^+^ T cell clones, was also reduced in siT6BP-treated cells (Fig 1D, left panel). Importantly, T6BP silencing did not interfere with the ability of HeLa-CIITA cells to present the cognate peptide recognized by Gag-specific T cells when the peptide was added exogenously (Fig 1D, right panel). These results suggest that T6BP silencing influences the generation of Gag-, Gag-LC3- and Gag-LC3_G120A_-derived endogenous epitopes and their subsequent presentation by MHC-II molecules to Gag-specific CD4^+^ T cells but does not affect the presentation of exogenous peptides by MHC-II molecules. Note that we obtained similar results with several siRNAs targeting different exons and introns of T6BP mRNA. We then sought to extend these observations to additional viral antigens. To this end, HeLa-CIITA cells treated with siCTRL or siT6BP were transfected with a plasmid encoding the immunodominant pp65 HCMV antigen and co-cultured with a pp65-specific CD4^+^ T cell line (Fig 1E). The viability and the percentage of pp65^+^ HeLa-CIITA cells were similar in both conditions (not shown). Remarkably, T6BP silencing also led to a strong reduction of CD4^+^ T cell activation (Fig 1E, left panel). As previously, the capacity of HeLa-CIITA cells to present the cognate pp65-derived peptide, added exogenously, was not affected (Fig 1E, right panel). These results demonstrate that regardless of the antigen tested, T6BP silencing dramatically influences the capacity of APCs to activate antigen-specific CD4^+^ T cells. The effect of T6BP silencing is broader than we initially anticipated as it impacts both autophagy-dependent and - independent endogenous viral antigen processing and presentation by MHC-II molecules.

### T6BP silencing lowers the stability of newly formed MHC-peptide complexes but does not significantly influence MHC-II molecule cell-surface expression and internalization

Although T6BP silencing does not significantly alter the capacity of cells loaded with exogenous peptides to activate antigen-specific CD4^+^ T cells (Fig 1D and E right panels), we asked whether T6BP might influence the internalisation of MHC-II molecules. Using flow cytometry, we first monitored on T6BP-silenced HeLa-CIITA cells, the expression levels of HLA-DR molecules using the L243 and TÜ36 antibodies that recognize mature HLA αβ and immature HLA αβ heterodimers associated with the invariant chain (Ii), respectively (Fig 2A, left panel: L243 and right panel: TÜ36). Using both antibodies, we noticed a slight increase of HLA-DR cell-surface expression levels on cells silenced for T6BP expression (Fig 2A). We next analysed the effect of T6BP silencing on the internalization kinetics of MHC-II molecules. HeLa-CIITA cells treated with siCTRL or siT6BP were coated at 4°C for 30 min with the L243 antibody that has been shown to act as an agonist of MHC-II molecules leading to their cellular internalization (De Gassart *et al*, 2008). The cells were then maintained at 4°C to monitor the antibody drop-off (Fig 2C) or incubated at 37°C to follow internalization of MHC-II molecules (Fig 3B). The cells were collected at the indicated time points and stained at 4°C with a labelled secondary antibody to detect the remaining L243 antibody conjugated with MHC-II molecules at the cell surface (Fig 2B and C). At 4°C, the mean fluorescent intensity (MFI) of HLA-DR molecules remained stable (Fig 2C). In contrast, at 37°C, the MFI dropped reaching a plateau after 40 minutes in both experimental conditions (Fig 2B, left panel), thus suggesting that the L243 antibody induced the internalization of HLA-DR molecules in the presence or absence of T6BP expression. As previously, compared to control condition, the silencing of T6BP slightly increased the expression levels (MFI) of HLA-DR molecules on the cell surface. However, it did not influence the kinetics of internalization of HLA-DR (Fig 2B, right panel).

**Figure 2.**
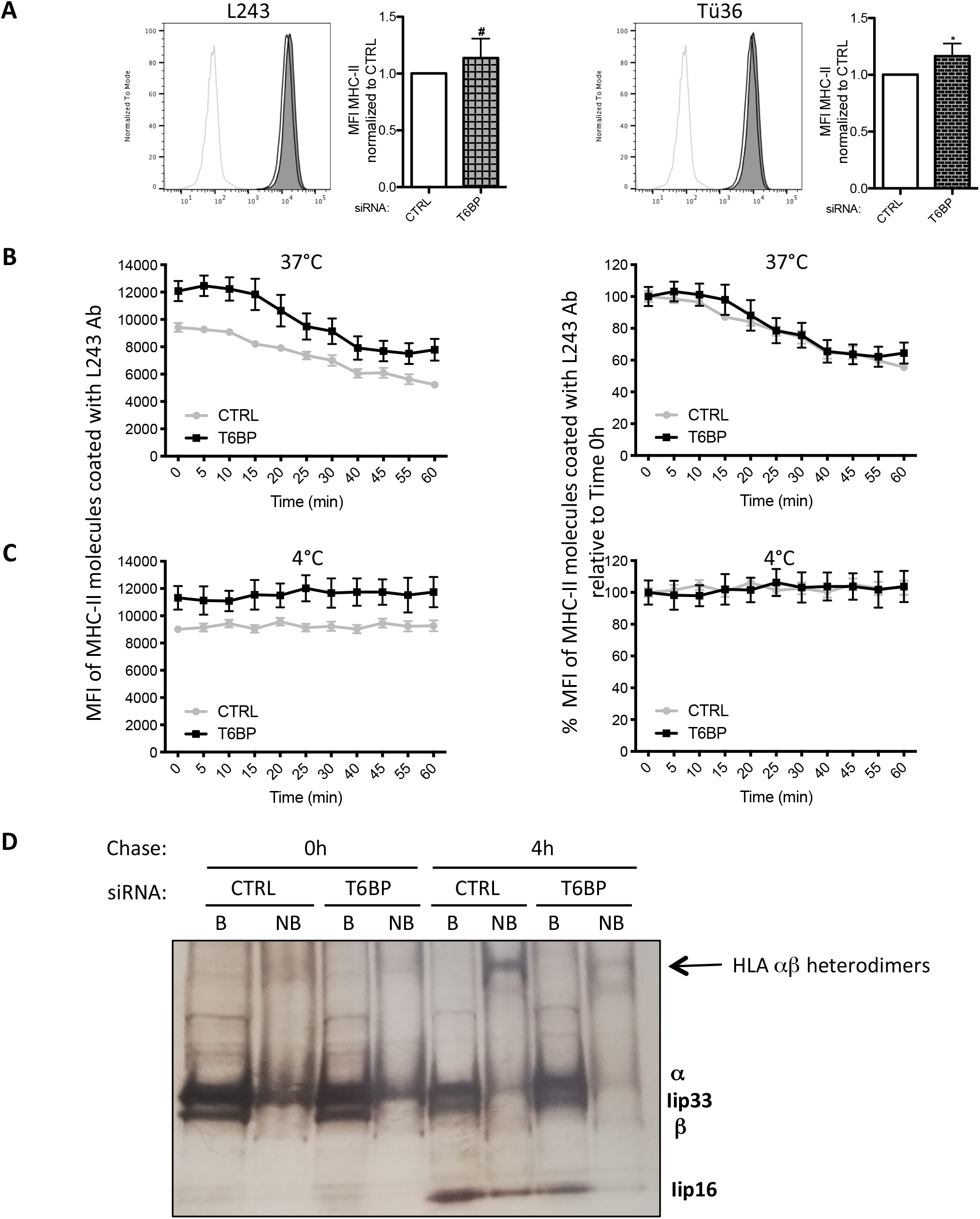
T6BP silencing influences the stability of HLA αβ dimers, mildly their cell-surface expression levels but has no impact on their internalization kinetics. **(A) Cell-surface expression of MHC-II molecules** was assessed using flow cytometry. HeLa-CIITA cells were transfected with siCTRL and siT6BP. 48h post-treatment, HLA-DR molecules were detected using L243 (left panels) and Tü36 antibodies (right panels) that recognize mature (HLA αβ heterodimers) and both mature and immature (HLA αβ and αβli complexes), respectively. Left, MFI of one representative experiment is presented as histogram. Right, at least five independent experiments were combined and presented as the means (+/-SD) of mean fluorescent intensity (MFI) standardized to the control conditions. Light grey lines: isotype negative controls; black lines and filled grey lines: anti-MHC-II staining of siCTRL- and siT6BP-treated cells, respectively. **(B)** and **(C) internalization kinetics of cell-surface HLA-DR molecules**. HeLa-CIITA cells were transfected as above. 48h post-treatment, mature HLA-DR molecules were stained at 4°C using L243 antibody. Cells were then incubated at 37°C (B) or at 4°C (C). At indicated time-points, cells were stained with a fluorescent secondary antibody at 4°C. Results are represented as MFI of MHC-II molecules stained with the L243 antibody (Ab) remaining at the cell surface (B and C, left panels), or as percentage (%) of MFI relative to time 0h (100%) (B and C, right panels). Results are representative of three independent experiments. CTRL: control. Mann-Whitney’s tests; *p<0.05; #p>0.05. **(D) T6BP silencing affects the formation of stable MHC-peptide complexes**. HeLa-CIITA cells were transfected as above and pulse-labeled with 35S-Met/Cys for 30 min, washed and chased for 4h at 37°C. MHC-II molecules were then immunoprecipitated using TÜ36 antibody and analyzed on SDS-PAGE after incubation of the immunoprecipitated protein complexes with SDS at 95°C (B: boiled) or room temperature (NB: non-boiled) to visualize α, β and Ii chains and SDS resistant αβ dimers, respectively. The bands corresponding to α, β and Ii chains are indicated. The arrow indicates the SDS-resistant stable HLA αβ heterodimers. This gel is representative of 2 independent experiments.

**Figure 3:**
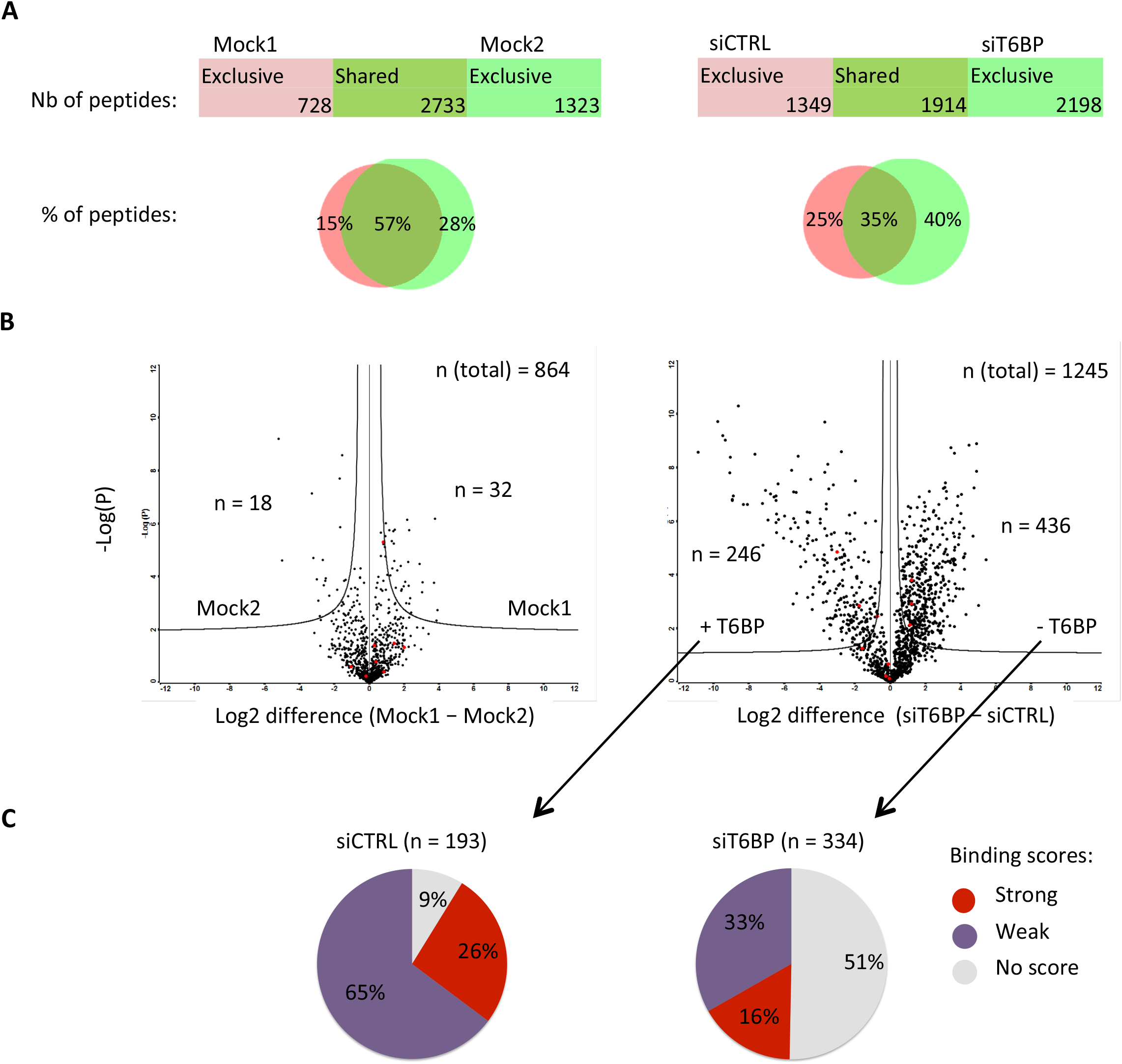
T6BP silencing alters the immunopeptidome of MHC-II molecules. **(A) Left panel**, mock-treated HeLa-CIITA cells were split and culture for 48h (giving rise to Mock1 and Mock2), then cells were lysed, MHC-II molecules were immunoprecipitated using TÜ39 antibody and the peptide-ligands sequenced using mass-spectrometry (LC-MS/MS). (A) **Right panel**, HeLa-CIITA cells were transfected with siCTRL and siT6BP siRNA and were treated as in the left panel. The number and the percentage among sequenced peptides (Venn diagrams) of exclusive or shared peptides for each condition are presented. **Quantitative (B) and qualitative (C) assessment of T6BP influence on the immunopeptidome**. Data from (A) were submitted to PLAtEAU algorithm to allow identification and label-free quantification of shared consensus core epitopes. **(B)** Volcano plots showing the log_2_ fold-change of core epitope intensity between siCTRL and siT6BP-treated cells (right panel) and Mock1 and Mock2 (left panel). For peptides exclusive to one or the other conditions, a background score was imputed to allow log2 fold-change presentation. An FDR of 0.01 and an S0 of 0.2 as correction factor for differences in the means were used. The resulting interval of confidence are highlighted by solid lines shown in each graph. The total number (n) of core epitopes and the number of epitopes with significant fold-change are indicated. **(C)** Relative binding affinities, presented as pie charts, of exclusive core epitopes identified by PLAtEAU in siCTRL (left) and siT6BP (right) conditions (number of epitopes are indicated in brackets). NetMHCIIpan was used to predict the relative affinities to HLA-DRb1*0102 expressed by HeLa-CIITA cells. The results are presented as stated from NetMHCIIpan analysis as Strong (for strong binders), Weak (for weak binders), and No score (for epitopes for which no binding score could be determined). Except for the Mock conditions, one representative experiment is shown out of two biological replicates. For each experiment, 5 technical replicates per sample were analyzed. Nb : number; %: percentage.

We then asked whether the silencing of T6BP might affect the stability of peptide-loaded HLA αβ heterodimers. A fraction of peptide-loaded MHC-II αβ heterodimers adopts a stable conformation resistant to dissociation by SDS at room temperature but not at 95°C (Germain & Hendrix, 1991). 48h post-transfection, we thus analysed the influence of siT6BP on the formation of SDS-stable HLA-DR αβ heterodimers. To this end, cells were pulsed with S^35^ labelled methionine and cysteine, and then chased for 4h, lysed and submitted to immunoprecipitation using TÜ36 that recognises HLA αβ heterodimers and HLA αβIi complexes but not free α, β and Ii chains (Benaroch *et al*, 1995). Prior loading onto the SDS-PAGE gel, samples were either boiled or incubated at room temperature for 30 minutes in SDS sample buffer (Fig 2D). Immediately after the pulse, boiling the samples revealed the α, β chains and the fragment Iip33/Iip35 of the invariant chain (Fig 2D, boiled, chase 0h). As expected from previous work, less α, β and Ii were detectable after 4h of chase (Fig 2D, non boiled, chase 4h) (Benaroch *et al*., 1995). The non-boiled samples revealed the existence of SDS-resistant stable peptide-loaded HLA-DR αβ heterodimers but only after 4h of chase (Fig 2D, non-boiled). Remarkably, in cells silenced for T6BP expression the band corresponding to SDS-resistant MHC αβ heterodimers was greatly reduced as compared to cells transfected with siCTRL (Fig 2D, non-boiled, chase 4h), strongly suggesting that T6BP expression facilitates the formation of stable peptide-loaded HLA molecules. Overall, these results show that T6BP silencing does not influence in a significant manner the cellular expression levels and the internalisation of MHC-II molecules. However, it has a significant impact on the stability of peptide-loaded MHC-II αβ heterodimers.

### T6BP silencing dramatically alters the immunopeptidome of MHC-II molecules

Various parameters affect the quality of peptide-loaded MHC-complexes, including the nature of the peptide itself (Roche & Furuta, 2015). We thus decided to analyse the global peptide repertoire (immunopeptidome) presented by MHC-II molecules. In addition, analysing the immunopeptidome might also reveal whether T6BP silencing affects a broader range of potential antigens. To study the immunopeptidome, HeLa-CIITA cells were either Mock-treated or transfected with CTRL or T6BP-silencing siRNA, MHC-II molecules were immunoprecipitated by using the TÜ39 antibody (specific to mature HLA-DP, DQ, and DR), and finally the peptide-ligands were identified using mass-spectrometry (LC-MS/MS). To assess the intrinsic variability of the MHC-II ligandome in HeLa-CIITA cells, we analysed simultaneously the ligandome of two samples from mock treated HeLa-CIITA cells that were split 48h prior to lysis and MHC-II immunoprecipitations (IP). In these settings, 57% of identified peptides were shared by both samples of mock-treated cells (Fig 3A, left panel). Two biological replicates of siRNA treated cells were analysed, one is presented in Fig 2. We observed that 1349 peptides (representing 25% of the peptides) were presented by MHC-II molecules exclusively in HeLa-CIITA cells expressing T6BP (Fig 3A, right panel, siCTRL). Remarkably, in the absence of T6BP expression, 2198 new MHC-II ligands (40% of the peptides) were identified (Fig 3A, right panel, siT6BP) and only 35% of the peptides (1914 peptides) were shared between control and the T6BP-silenced conditions (Fig 3A, right panel). Overall, siT6BP treated cells shared significantly less MHC-II peptide ligands with the three other experimental conditions (Mock1, Mock2 or siCTRL) (Fig S2A). Together these results show that, although there is an intrinsic variability of the MHC-II ligandome in cells, the absence of T6BP has a pronounced and dramatic influence on the repertoire of peptides presented by MHC-II molecules.

We then asked whether T6BP expression might affect the source of peptides, meaning the set of proteins supplying peptides for MHC-II loading. To this end, we submitted the LC-MS/MS data to cell component enrichment analysis using Funrich software (Pathan *et al*, 2015). As expected from previous work (Dengjel *et al*., 2005; Marcu *et al*, 2021), according to Funrich annotation and as compared with the human proteome, in both siCTRL and siT6BP treated cells, the MHC-II-ligand source proteins were enriched in cellular fractions belonging to membranes, secretory pathways (e.g. exosomes and vesicles) but also to the cytosolic compartment of the cell (Fig S2B). In siCTRL treated cells MHC-II source proteins were also augmented in nuclear fractions, which was not the case for siT6BP-transfected cells. In contrast, MHC-II-ligand source proteins belonging to the cytoplasm were enriched in siT6BP-treated cells (Fig S2B). Overall, this analysis shows that the bulk of MHC-II-ligand source proteins is provided by the same cellular fractions mainly membranes, secretory pathways and the cytosol. However, the presence of T6BP might also favour the presentation of peptides derived from the nucleus.

We then sought to analyse the influence of T6BP on the relative abundance and the quality, meaning the affinity to MHC-II molecules, of peptides presented. However, peptides eluted from MHC-II molecules are variable in length and one core epitope required for MHC-II binding, usually 13 amino acid long, can be found in multiple peptides with N- and C-terminal extensions (Rammensee *et al*, 1999). To circumvent this limitation, we adapted a protocol, published by Alvaro-Benito *et al*., to identify the core epitopes within our data sets using the Peptide Lansdscape Antigenic Epitope Alignment Utility (PLAtEAU) algorithm (Alvaro-Benito *et al*, 2018). A total of 864 and 1245 unique core epitopes were identified in the groups Mock1/Mock2 and siCTRL/siT6BP, respectively. PLAtEAU also allows calculating of the relative abundance of the core epitopes based on the LC-MS/MS intensities of peptides containing the same core epitope. Between the two mock-treated samples, around 5% of the core epitopes showed a significant difference in their relative abundances (Fig 3B, left panel). Remarkably, between the siCTRL and siT6BP conditions, 55% of the core epitopes displayed a relative abundance that was significantly different between siCTRL and siT6BP conditions, with 436 and 246 epitopes more abundant in the siT6BP or siCTRL-treated cells, respectively, among 1245 peptides (Fig 3B, right panel). Therefore, T6BP silencing influences both the peptide repertoire (Fig 3A) and the relative abundance of a majority of the core epitopes presented by MHC-II molecules (Fig 3B).

Having identified the core epitopes, using NetMHCIIpan4.0 algorithm (Reynisson *et al*, 2020), we next analysed the relative binding affinities of exclusive peptides, identified in siCTRL and siT6BP conditions, to the HLA-DRβ1*0102 allele that is expressed by HeLa-CIITA cells. Strikingly, when T6BP is expressed (siCTRL), more than 90 % of the epitopes were predicted to be HLA-DRβ1*0102 binders (26% and 65%, strong and weak binders, respectively). In contrast, in T6BP-silenced cells, below 50% of peptides were predicted to bind HLA-DRβ1*0102 (16% and 33%, strong and weak binders respectively). We also compared the predicted binding capacities of core epitopes exclusive to the Mock1 or Mock2 conditions and it did not reveal a significant difference in terms of relative affinity (not shown). Together, these results suggest that in the absence of T6BP, the peptides presented by HLA-DRβ1*0102 molecules have a predicted weaker relative affinity.

Note that we also analysed on the same samples whether T6BP might influence the immunopeptidome of MHC-I molecules. Cells transfected with siCTRL or siT6BP and the two mock-treated samples were submitted to IP but using the pan anti-MHC-I antibody, W632, and the peptide ligands sequenced using LC-MS/MS. We did not observe a significant influence of T6BP on the percentage of shared or exclusive peptides comparing the two mock-treated samples and the siCTRL/siT6BP conditions (Fig S3A and not shown). We further analysed the relative affinities of the exclusive peptides bound to siCTRL- and to siT6BP-treated cells. In contrast to MHC-II ligands, MHC-I molecules present short 9-mer peptides that correspond to the core epitope (Rammensee *et al*., 1999). We thus directly analysed the relative affinities using the NetMHCpan 4.0 algorithm (Jurtz *et al*, 2017). The percentages of strong, weak and non-binder peptides were similar in control and T6BP-silenced cells (Fig S3B). Therefore, the action of T6BP seems to be limited to the MHC-II-restricted antigen presentation pathway. T6BP expression has a strong influence on the peptide repertoire, the relative abundance and the relative affinity of epitopes presented by MHC-II molecules.

### T6BP silencing affects the cellular localization of the MIIC

We then asked whether T6BP might play a role in the intracellular trafficking of MHC-II molecules, before mature peptide-loaded MHC-II molecules reach the plasma membrane. To this end, using confocal microscopy analysis, we evaluated in HeLa-CIITA cells the effect of T6BP silencing on endo-lysosomal compartments. We first confirmed previous results (Petkova *et al*, 2017) showing that T6BP silencing leads to the accumulation of LC3-positive puncta corresponding to autophagosomes (Fig S4A-B). Note that at steady state in HeLa-CIITA-cells, T6BP did not co-localize with LC3^+^ puncta (Fig S4B). We extended this observation using electron microscopy of the morphology of siRNA-treated cells and confirmed that large (around 1 μm in length) vesicles with a double membrane (with 20 to 30 nm interspace) accumulated in about 80% of the cells silenced for T6BP expression (Fig S4E). These structures where not observed in the siCTRL-treated cells. Two independent experiments were performed and at least 40 cells for each treatment were analysed. We then analysed, by confocal microscopy, the subcellular localization of MHC-II molecules together with markers of autophagosomes, late endosomes and lysosomes. In HeLa-CIITA cells (Bania *et al*, 2003), MHC-II molecules, stained with an antibody to mature HLA-DR molecules (L243), localized in patches corresponding to intracellular vesicles that did not include T6BP (Fig 4A). Whatever the siRNA treatment, MHC-II molecules did not co-localize with LC3-positive puncta (Fig S4C and D). However, when compared to the control condition, MHC-II molecules seemed to localize closer to the nucleus upon T6BP silencing (Fig 4A). We determined the average distance to the nucleus and the number of vesicles in more than 150 cells representing over 20 000 MHC^+^ puncta (Fig 4B). We noticed a slight but significant decrease of MHC-II-positive vesicle distance to the nucleus upon T6BP silencing (Fig 4B). The numbers of MHC-II^+^ spots per cell were not significantly different (Fig 4B). Using an anti-LAMP1 antibody, we then analysed effect of T6BP expression on late endo-lysosomal compartments. As compared to control cells, we observed a significant re-localization of LAMP-1-positive vesicles at the proximity of the nucleus in T6BP-silenced cells (Fig 4C and D). The number of LAMP-1-positive vesicles was globally unchanged (Fig 4C and D).

**Figure 4.**
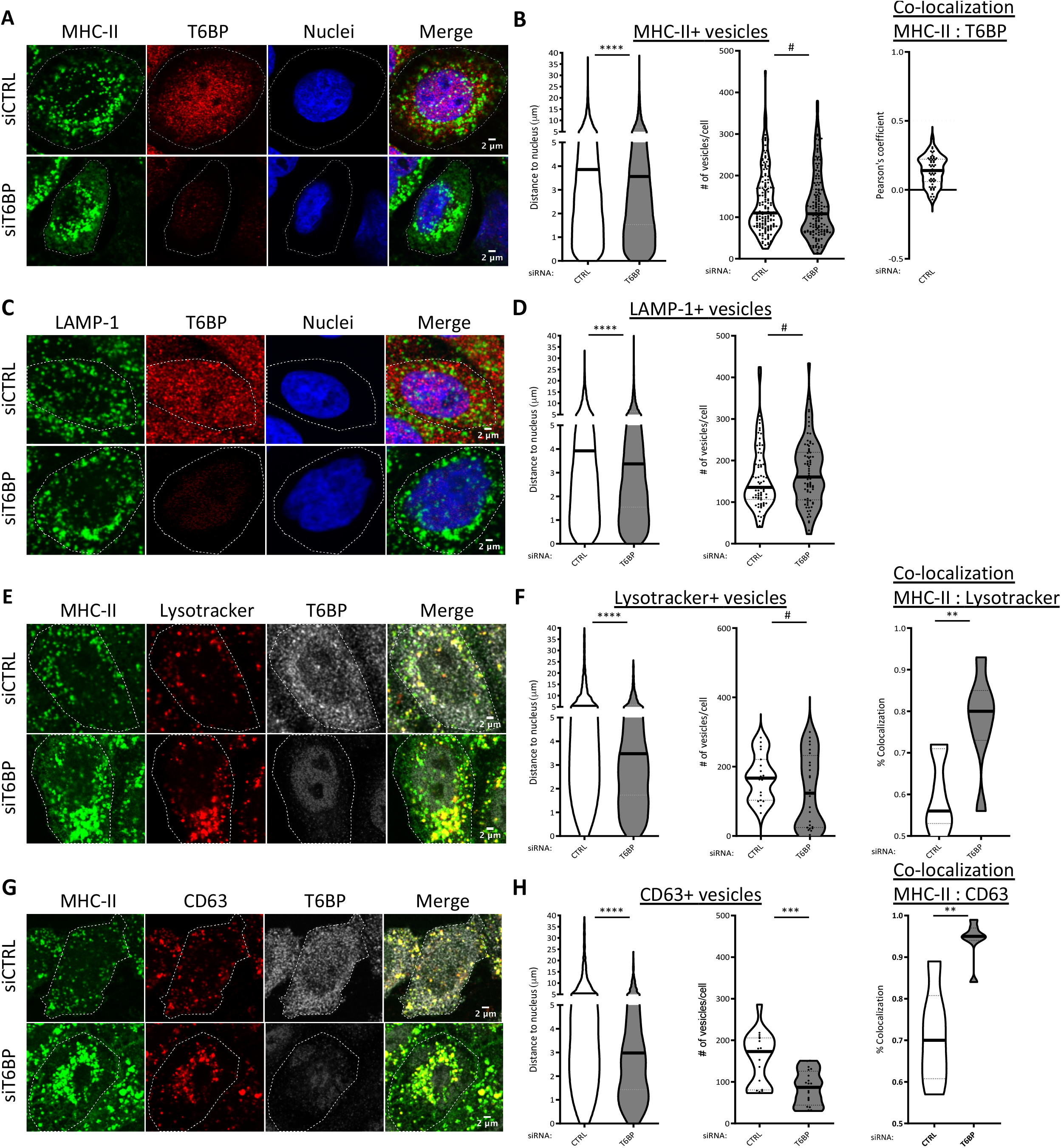
T6BP silencing leads to perinuclear relocalization of the MIIC. **(A)** MHC-II and T6BP expressions were assessed using confocal microscopy. HeLa-CIITA cells were transfected with control and T6BP-silencing siRNAs. 48h post-treatment, MHC-II and T6BP were detected using L243 and anti-T6BP antibodies, respectively, and revealed with species specific secondary antibodies. Nuclei were stained using DAPI. **(B)** Quantitative analysis using in-house ImageJ script displaying distance of each MHC-II+ vesicles to the nucleus and number of vesicles per cell. At least 20,000 vesicles from 160 cells corresponding to 5 independent experiments were analyzed. Right panel, quantification in the siCTRL cells of the colocalization between MHC-II+ and T6BP+ dots using Pearson’s coefficient where the dotted lines (at 0.5) indicate the limit under which no significant co-localization is measured (number of cells = 54). **(C)** As in A, T6BP and LAMP-1 expressions were analyzed. **(D)** The localization of LAMP-1+ vesicles and the number of vesicles per cell were quantified as in B. At least 2000 vesicles from 20 cells corresponding to 2 independent experiments were analyzed. **(E)** As in A, adding Lysotracker staining. **(F)** As in B, quantitative analysis of Lysotracker+ vesicles: localization to the nucleus and number of vesicles per cell. Colocalization of Lysotracker+ vesicles with MHC-II+ puncta was analyzed using JACoP plugin (scales start at 0.5 above which the % of co-localization is considered significant), number of vesicles > 2000 from at least 2 independent experiments were analyzed corresponding to 20 cells. **(G)** As in E, CD63 and MHC-II expressions were assessed. **(H)** Quantitative analysis as represented in F. At least 2000 vesicles from 20 cells corresponding to 2 independent experiments were analyzed. In graphs representing the number of vesicles per cells, each dot displayed corresponds to a single cell. Scale bars, 2μm. CTRL: control. Mann-Whitney’s tests; *p<0.05; **p<0.002; ***p<0.0003; ****p<0.0001; #p>0.05.

We next asked whether the MIIC itself might be affected by T6BP silencing in HeLa-CIITA cells. The MIIC is a labile ill-defined acidified compartment that has been shown to be positive for multiple markers that do not always overlap (Roche & Furuta, 2015). We used Lysotracker that stains acidified compartments, CD63, a tetraspanin molecule anchored to the membrane of the intraluminal vesicles of the MIIC (Roche & Furuta, 2015), HLA-DM, the chaperone involved in MHC-II peptide loading, and MHC-II molecule staining to identify the MIIC. In the absence of T6BP, we observed a pronounced, statistically significant, re-localization of Lysotracker-positive vesicles close to the nuclei but the number of Lysotracker-positive vesicles was unchanged (Fig 4E and F). Remarkably, in siT6BP-treated cells, Lysotracker-positive vesicles showed increased co-localization with MHC-II molecules (Fig 4E and F). Upon T6BP silencing, CD63-positive vesicles were also strongly re-localized around the nucleus and showed an increased co-localization with MHC-II-positive puncta (Fig 4G and H). The number of CD63-positive vesicles was strongly reduced in T6BP-silenced cells (Fig 4H). Finally, we analysed the influence of T6BP silencing on the cellular localization of HLA-DM. As for CD63, in T6BP-silenced cells, HLA-DM^+^ vesicles were strongly re-localized around the nucleus and their number was reduced as compared to mock-treated cells (Fig S3F and G). In summary, in T6BP-silenced cells, LAMP-1^+^, HLA-DM^+^, CD63^+^ MHC-II^+^ and Lysotracker^+^ MHC-II^+^ vesicles showed a repositioning at the proximity of the nucleus (Fig 4).

siT6BP silencing also induced a slight accumulation of EEA1+ early endosomes in the perinuclear region (Fig S5A and B). Whatever the treatment, no colocalisation between EEA1 and MHC-II was observed (Fig S5A and B). Also note that overall, T6BP-silencing did not significantly reduce the expression levels of the various markers analysed using flow cytometry analysis (Fig 2), WB (Fig 4 and 5) and IF (Fig S5C-H). Taken together these results show that T6BP expression influences the positioning of the MIIC, which is re-localized closer to the nucleus in the absence of T6BP. This repositioning of the MIIC could at least partially account for the defect of MIIC maturation and could sustain the dramatic changes of the global MHC-II peptide repertoire, observed in the absence of T6BP.

**Figure 5.**
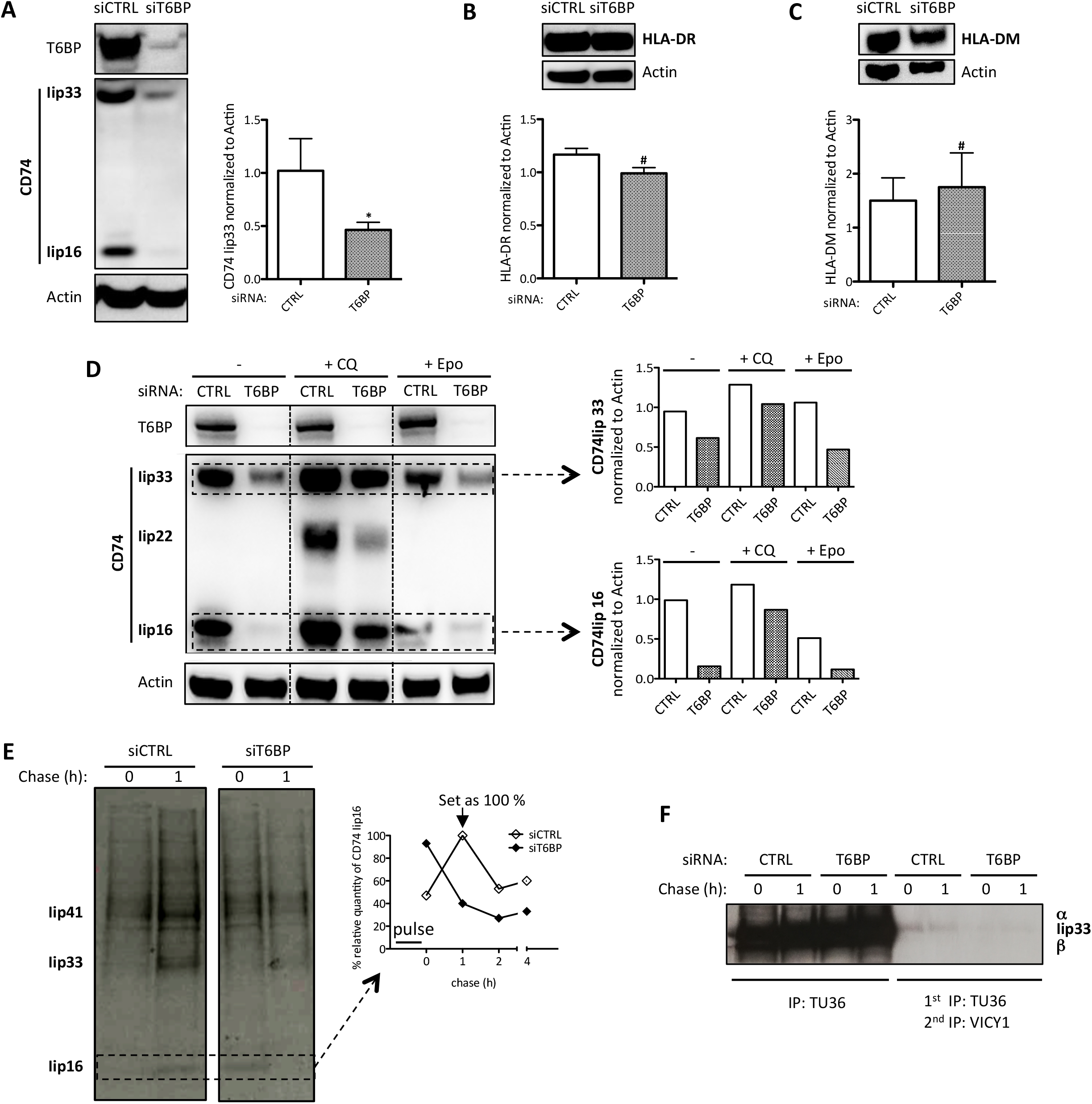
T6BP silencing leads to exacerbated CD74 (Ii) degradation. **(A)** CD74 expression was assessed using Western Blot. HeLa-CIITA cells were transfected with siCTRL and siT6BP. 48h post treatment, the Iip33 CD74 isoform, a degradation product (Iip16), T6BP and actin were detected using indicated antibodies. Left panel, a representative Western Blot experiment is shown. Right panel, expressions levels were quantified using ImageJ and are presented as ratios of CD74 to actin used as control housekeeping gene expression. **(B)** As in (A) with the same samples but assessing HLA-DR and actin expression levels on a different blot. Top panel, a representative Western Blot experiment is shown. Bottom panel, HLA-DR expression was quantified using ImageJ and presented as a ratio to actin. **(C)** As in (B) assessing HLA-DM and actin expression levels. Results are representative of at least three independent experiments. **(D)** CD74 expression is partially recovered by blocking lysosomal acidification. As in (A), following siRNA transfection of Hela-CIITA cells, CD74 expression was assessed. The last 16h prior to harvesting, cells were treated with chloroquine (CQ) or epoxomicin (Epo). Left panel, membranes were blotted using anti-T6BP, -CD74 and –actin antibodies. The different degradation fragments of CD74 are indicated (Iip33, Iip22 and Iip16). Right panel, expression levels of Iip33 (top) and Iip16 (bottom) were quantified using ImageJ and are presented as ratios of each fragment to actin. These results are representative of at least three independent experiments. **(E and F)** Analysis of CD74 proteolysis. HeLa-CIITA cells were transfected with siCTRL and siT6BP. 48h post treatment, cells were pulsed for 30 min with ^35^S-Met/Cys, washed and chased for 1, 2 and 4h. HLA-DR/CD74 complexes were first immunoprecipitated using Tü36 antibody (**E**) and then reimmunoprecipitated with VICY1 antibody. (**F**). Samples were boiled and analyzed using SDS-PAGE. The bands corresponding to CD74 isoforms (Iip41 and Iip33) and the cleavage products (Iip16) are indicated. **(E right panel)** Iip16 expression was quantified using ImageJ and presented as a percentage of Iip16 normalized to the highest quantity of Iip16 detected after 1h of chase in the control condition. Results are representative of two independent experiments. Ii: invariant chain (CD74); CTRL: control. Wilcoxon’s tests; *:p<0.05; #:p>0.05.

### T6BP silencing leads to exacerbated CD74 degradation in HeLa-CIITA cells

CD74 (Ii) is essential for the traffic and the maturation of MHC-II molecules within the cell (Bakke & Dobberstein, 1990; Lotteau *et al*., 1990; Neefjes *et al*., 1990; Roche *et al*., 1991). In the MIIC, CD74 degradation is also strictly regulated to ensure appropriate loading of MHC-II molecules with high affinity peptides (Manoury *et al*., 2003; Nakagawa *et al*., 1998; Riese *et al*., 1996; Shi *et al*., 2000). We thus assessed the effect of T6BP silencing on CD74 expression. Note that in humans, among the four CD74 isoforms (Iip33, Iip35, Iip41 and Iip43) Iip33 is the most abundant (Thibodeau *et al*., 2019). As previously, HeLa-CIITA cells were transfected with siRNAs and CD74 expression was assessed by western blotting. In control cells, Iip33 was readily detected together with its cleavage product Iip16 (Fig 5A). In contrast, upon T6BP silencing, Iip33 and Iip16 detection was strongly decreased (Fig 5A). Normalised to the housekeeping gene (actin), the decrease of Iip33 expression reached up to 50% (Fig 5A, right panel). As control, using western blotting (WB), we also assessed the expression level of HLA-DR molecules. As expected from our flow cytometry and microscopy results (Fig 2 and 4), the global expression of HLA-DR molecules was similar in siCTRL- and siT6BP-treated cells (Fig 5B). In addition, we asked whether T6BP silencing might also affect the expression of HLA-DM, the chaperone involved in quality control of peptide loading on MHC-II molecules. As for HLA-DR, the expression levels of HLA-DM were not affected by the extinction of T6BP expression (Fig 5C). Using confocal microscopy, we confirmed that T6BP-silenced cells exhibit about 50% lower expression levels of CD74 than control cells (Fig S6A and B). Note that, in control cells, the T6BP staining did not co-localize with CD74 (Fig S6A and C).

We then asked at which level CD74 expression was affected, meaning at the RNA or protein levels. To this end, we first used RT-qPCR to monitor CD74 and T6BP mRNA relative quantities in control and T6BP-silenced HeLa-CIITA cells. As anticipated T6BP mRNA levels were strongly reduced in siT6BP-treated cells (Fig S6D). In contrast, mRNA levels of CD74 were not influenced by T6BP silencing (Fig S6E). We next analysed whether CD74 expression could be reversed, in siT6BP-silenced cells, upon treatment with drugs inhibiting lysosomal or proteasomal degradation, using Chloroquine or Epoxomycin, respectively. As expected, treatment of control cells with Chloroquine, leads to the accumulation of Iip33 and Iip16, and allowed the detection of the intermediate degradation fragment Iip22 (Fig 5D). Remarkably, when compared to control untreated cells, Chloroquine treatment allowed a strong recovery of both Iip33 and Iip16 expression in siT6BP-treated cells (Fig 5D). In contrast, Epoxomycin treatment did not affect the expression levels of CD74 (Fig 5D).

These results prompted us to analyse the kinetics of degradation of CD74 complexes. To address CD74 proteolysis in control and T6BP silenced Hela-CIITA cells, 48h post-transfection cells were pulsed with S35 Met/Cys for 30 min and chased for different times. HLA-CD74 complexes were isolated by immoprecipitation with TÜ36 antibody and in a second set of experiments CD74 was first immoprecipitated with TÜ36 antibody and reimmunoprecipitated using VICY1 antibody, which binds the cytosolic tail of CD74. As shown in Fig 5E, in control cells, CD74 isoforms (Iip41 and Iip33) and its fragments (Iip16) appeared in the pulse and during the chase. In contrast, when Hela-CIITA was silenced for T6BP much less Ip41, Iip33 and Iip16 were detected. In fact, in the control conditions, the invariant chain degradation product, Iip16, was detected mostly after 1h of chase (Fig 5E). In contrast, Iip16 was immediately detected after the pulse (time 0h) in the T6BP-silenced cells (Fig 5E) suggesting a rapid CD74 degradation (see quantification on the right panel). Furthermore, when HLA/CD74 complexes where first immunoprecipitated with TÜ36 antibody and reimmunoprecipitated with VICY1 antibody, much less CD74 (Ip33) was detected in T6BP silenced cells (Fig 5F). Altogether these results suggest a much faster degradation of CD74 either free or associated with MHC-II molecules in the absence of T6BP.

Interestingly, the expression of CD74 has been shown to influence both the cellular trafficking and the immunopeptidome of MHC-II molecules (Muntasell *et al*, 2004). It has also been suggested that in cells lacking MHC-II expression, CD74 might regulate endosomal maturation (Schröder, 2016).

### T6BP interactome reveals novel binding partners

To decipher the mechanism by which T6BP influences MHC-II-restricted endogenous antigen presentation, we decided to define the interactome of T6BP in HeLa-CIITA cells. To this end, HeLa-CIITA cells were transfected with a plasmid encoding GFP-T6BP, a construct that was previously functionally characterized (Morriswood *et al*., 2007), and as negative control a plasmid encoding GFP (Fig S7A). We then performed a large-scale immunoprecipitation (IP) using GFP as a tag. Three biological replicates were performed. As quality control, the lysates and the various fractions of the immunoprecipitation procedure were analysed using coomassie blue staining and anti-GFP staining in Western Blot (Fig S7B). The immunoprecipitation products of the three replicates were then submitted to mass spectrometry (LC-MS/MS) analysis to identify the proteins interacting with GFP-T6BP. (Fig S7C). The Uniprot *Homo sapiens* database was used to assign a protein name to the peptides sequenced by MS. The results were compared against the 3 IP-replicates of GFP transfected cells and a bank of 8 control experiments, also performed with GFP-Trap agarose magnetic beads, using the online contaminants database http://CRAPome.org (Mellacheruvu *et al*, 2013). Using a fold-change (FC) threshold of ≥ 10 and a Significance Analysis of INTeractome (SAINT) probability threshold of ≥ 0,8 (Choi *et al*, 2011), we identified 116 high-confidence T6BP proximal proteins (Fig 6 and Table S1). These included previously known T6BP-interactants such as the E3 ligase ITCH (Shembade *et al*, 2008), the kinase TBK1 (Richter *et al*, 2016) and TRAF2, all involved in NF-κB signalling pathways (Shembade *et al*, 2010). This screen also confirmed that T6BP interacts with the autophagy receptor p62 (SQSTM1) (Mildenberger *et al*, 2017). It revealed novel potential partners that might bind directly or as part of larger T6BP-associated protein complexes. These candidates could be grouped in 6 major functional cellular pathways based on Ingenuity (Fig 6). As expected, from the known T6BP functions, some candidate-partners were enriched in the Ubiquitine/Proteasome (p-value = 3,16E-18) and NF-κB signalling (p-value = 1,15E-02) pathways but also in the unfolded protein response (UPR)/protein folding (p-value = 2,69E-04), endocytosis (p-value = 6,03E-05), and antigen presentation (p-value = 2,34E-06) pathways (Fig 6). The antigen presentation group includes HLA-A, -C, -DQα1, -DRβ1 molecules (Fig 6). In the UPR/protein folding group, the ER-resident chaperone protein Calnexin (CANX) is 22 times enriched in T6BP-GFP IP as compared to GFP only IP (Fig 6 and Table S1). CANX drew our attention because: 1) it has been shown to interact with CD74 (Anderson & Cresswell, 1994) and 2) the inhibition of CANX/CD74 interaction induces CD74 degradation without influencing the formation of MHC-II complexes (Romagnoli & Germain, 1995).

**Figure 6.**
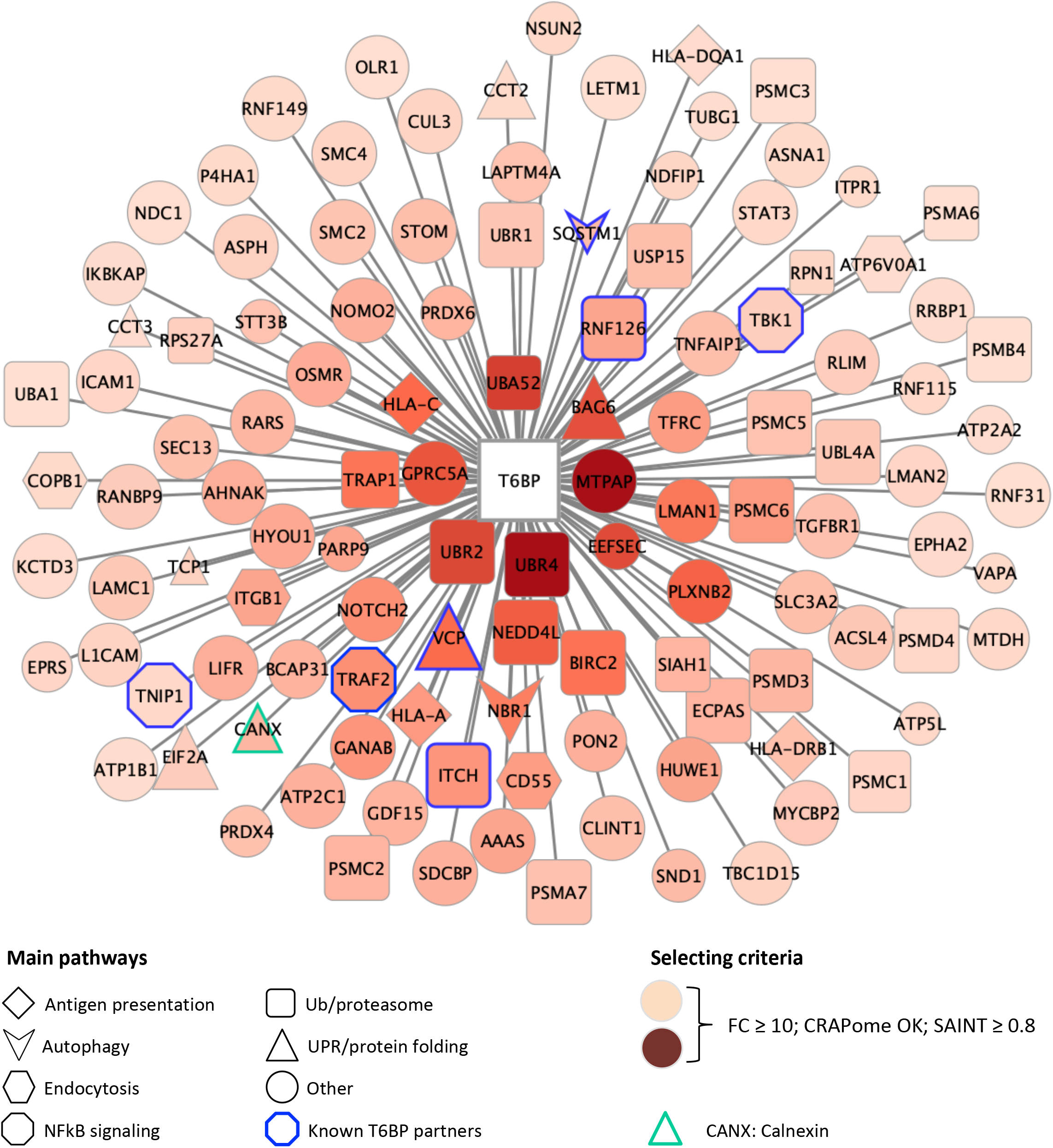
T6BP interactome reveals novel binding partners. Diagram of the T6BP protein interaction network identified by immunoprecipitation followed by LC-MS/MS and represented using Cytoscape. T6BP (white) coupled to GFP was immunoprecipitated using anti-GFP camel antibodies (GFP-Trap Chromotek). GFP alone was used as control. Proteins were considered as relevant partners based on the following criteria: at least 2 peptides were identified by LC-MS/MS, the fold-change (FC) to the GFP control condition ≥ 10, SAINT probability threshold ≥ 0,8. For data analysis the Resource for Evaluation of Protein Interaction Networks (REPRINT) and its contaminant repository (CRAPome V2.0) were used. The edge’s length is inversely proportional to the FC score (short edge = high FC) and the node’s color intensity is directly proportional to the FC score (the more intense the higher the FC). The size of the node is directly proportional to the SAINT score (lower confidence = smaller node). Blue border indicates previously described partners of T6BP. The shape of the node highlights the functional pathway in which the candidate protein is enriched based on Ingenuity. Green border indicates Calnexin = CANX.

### T6BP and Calnexin interaction is observed in professional APC and relies on Calnexin cytosolic tail expression

To verify that CANX interacts with T6BP, as previously, we immunoprecipitated GFP-T6BP and GFP from HeLa-CIITA transfected cells and analysed, using Western blot, the presence of Calnexin in the immunoprecipitated fractions. The IP of the GFP transfected cells lead to the immunoprecipitation of a larger GFP-positive fraction than the IP of the T6BP-GFP transfected cells (Fig S7D, anti-GFP Ab). Nonetheless, we observed a 3-fold enrichment of CANX in cells transfected with GFP-T6BP as compared to GFP-transfected cells (Fig S7D). In HeLa-CIITA cells, we further confirmed the specific interaction between T6BP and CANX using two different techniques: 1) by immunoprecipitation of endogenous T6BP and endogenous CANX, we revealed that the IP fractions of anti-T6BP and anti-CANX IPs contained CANX and T6BP, respectively (Fig 7A-B); 2) by proximity ligation assay (PLA) on control cells and siT6BP-transfected cells a specific T6BP/CANX interaction was detected only in cells expressing T6BP (Fig 7C). To demonstrate that T6BP/CANX interaction is not restricted to Hela-CIITA cells, we then immunoprecipitated endogenous T6PB and CANX from two professional APCs, namely B cells and DC. As in Hela-CIITA, we revealed in the anti-T6BP and anti-CANX IP fractions, of both B cells and DC, the presence of CANX and T6BP, respectively (Fig 7A-B). All together, these results strongly support that CANX interacts with T6BP directly or as part of a larger protein complex.

**Figure 7.**
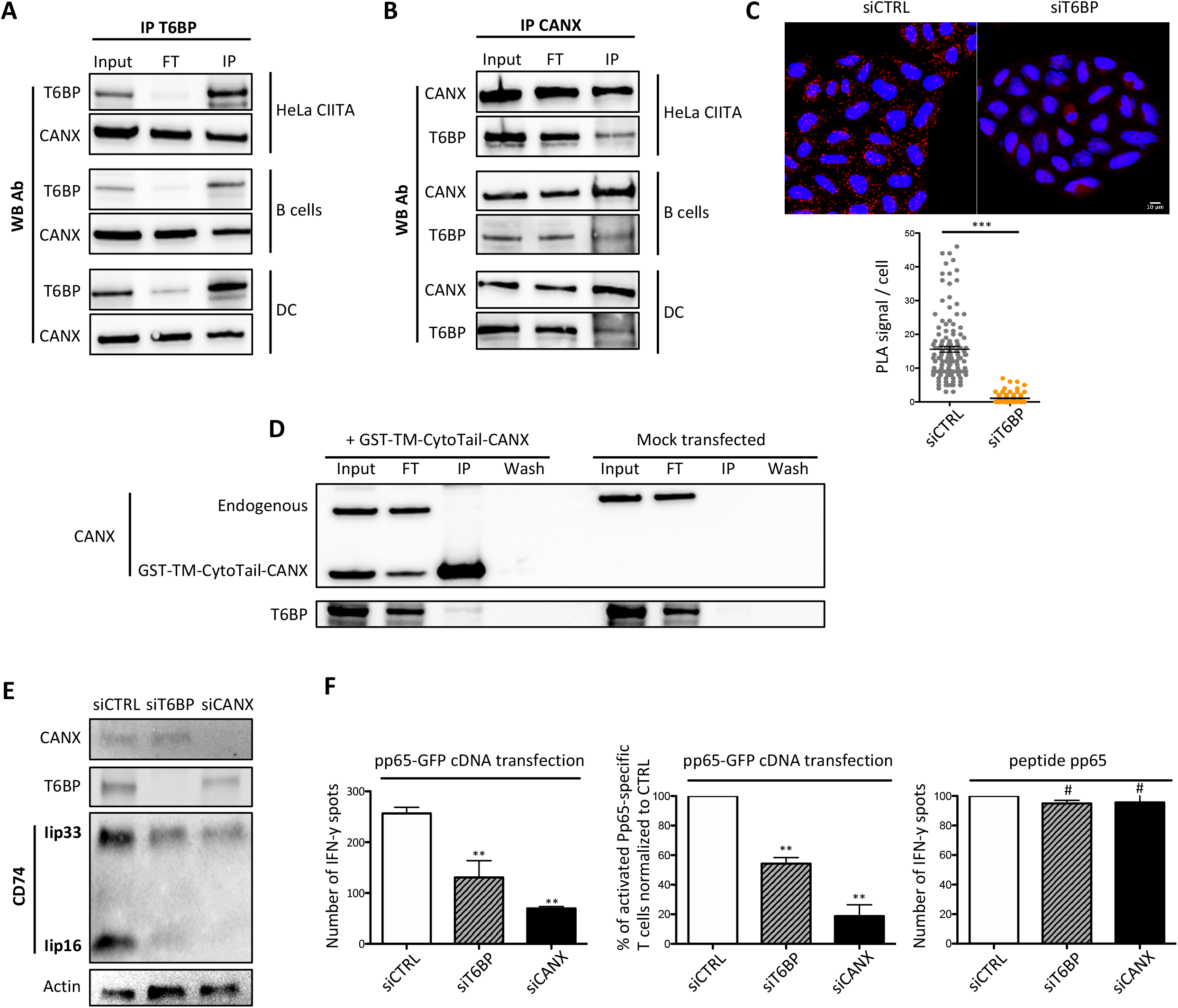
Calnexin (CANX) interacts with T6BP through its cytosolic tail and stabilizes CD74 thus favouring MHC-II-restricted presentation to CD4^+^ T cells. **(A-B) CANX co-immunoprecipitates with T6BP in model and professional APCs**. Endogenous T6BP was immunoprecipitated from HeLa-CIITA, B (DG75) and dendritic-like (KG-1, DC) cells. The input, the flow through (FT) and the immunoprecipitation (IP) fractions were analyzed by Western blot using the indicated antibodies. **(B)** As in (A) but using anti-CANX Ab for the IP. For A and B, one representative experiment out of 3 independent experiments is shown. **(C) CANX interacts with T6BP using proximity ligation assay (PLA)**. 48h post-transfection with the indicated siRNAs, HeLa-CIITA cells were fixed, stained with anti-T6BP and anti-CANX antibodies and proximity revealed using PLA (Duolink). Nuclei were stained using DAPI. Top panel, two representative fields are shown out of three 3 independent experiments. Bottom panel, quantitative analysis using ImageJ displaying the number of PLA per cell. PLA were quantified in at least 130 cells corresponding to 3 independent experiments. Scale bars, 10μm. CTRL: control; Mann-Whitney’s tests; ***: p<0.0003. **(D) Calnexin interacts with T6BP through its cytosolic tail**. HeLa-CIITA cells were transfected with a plasmid encoding the GST-tagged transmembrane (TM) and cytosolic (CytoTail) domains of CANX (GST-TM-CytoTail-CANX) or mock treated and immunoprecipitated with anti-GST antibodies. The input, FT, the wash and the IP fractions were analyzed by Western blot and revealed using the indicated antibodies. One representative experiment out of 3 independent experiments is shown. **(E) Silencing of T6BP does not influence CANX expression levels while silencing of CANX reduces the level of CD74 expression**. HeLa-CIITA cells transfected with the indicated siRNAs and samples analyzed, 48h post transfection, by Western blot with the indicated antibodies. These western blot results are representative of at least 3 independent experiments. **(F) CANX silencing dampens MHC-II-restricted antigen presentation to CD4**^**+**^ **T cells**. HeLa-CIITA cells were treated with the indicated siRNAs and transfected with a plasmid encoding pp65 HCMV antigen fused to GFP. HeLa-CIITA cells were then co-cultured with pp65-specific T cells. Left panel, a representative experiment is shown. T6BP and CANX silencing was confirmed using WB (not shown). Middle panel, three independent experiments are combined and presented as the mean percentage (+/-SD) of activated cells producing IFN© normalised to CTRL conditions. Right panel, influence of T6BP silencing on peptide presentation by Hela-CIITA cells. The cognate peptide was added exogenously (pp65, 0,5μg/mL) on siRNA-treated cells (2h, 37°C), washed and T cell activation monitored using IFN©-ELISPOT. The background secretions of IFN© by CD4^+^ T cells co-cultured with mock-treated HeLa-CIITA cells were used as negative controls and subtracted. CTRL: control. Wilcoxon’s tests; **p<0.01; #p>0.05.

In order to characterize the molecular interactions between CANX and T6BP, we then engineered a construct containing solely the transmembrane domain and the cytosolic tail of CANX fused to a GST tag. Since T6BP expression is mainly cytosolic, we reasoned that T6BP and CANX might interact through the cytosolic tail of CANX. The GST-TM-cytosolic Tail construct was transfected in Hela-CIITA cells and immunoprecipitated using anti-GST antibodies. Western blotting revealed that T6BP is co-immunoprecipitated with anti-GST in cells transfected with GST-TM-cytoTail and not in mock-transfected cells (Fig 7D). Taken together, we demonstrate here that T6BP interacts with CANX through its cytosolic tail. We propose that, through the binding to the cytosolic tail of CANX, T6BP promotes the interaction of CANX with CD74 leading to CD74 stabilization.

### Calnexin silencing induces CD74 degradation and reduces MHC-II-restricted presentation to CD4^+^ T cells

Finally, we investigated a direct role of CANX in MHC-II-restricted antigen presentation. To this end, we screened several siRNA targeting Calnexin and identified an siRNA whose transfection led to a strong decrease of CANX expression (siCANX) (Fig S7E), without affecting neither HLA-DR and -DM expression levels nor the viability of transfected Hela-CIITA cells (not shown). Using Western blot, we first analysed the influence of CANX-silencing on CD74 expression. In siCANX-treated cells, we observed a strong reduction of CD74 expression levels (Fig 7E) reminiscent of what we observed in cells silenced for T6BP expression (Fig 5A and 7E). Note that neither siT6BP nor siCANX transfections lead to an increase expression of CANX or T6BP respectively (Fig 7E and S7E). To monitor the influence of Calnexin on antigen presentation, HeLa-CIITA cells were transfected with siCANX side by side with siCTRL or siT6BP used as negative and positive controls, respectively. As in Fig 1E, the cells were then transfected with the plasmid encoding the HCMV pp65 antigen fused to GFP. The levels of pp65-GFP expression were monitored using flow cytometry (not shown) and the cells were co-cultured with the anti-pp65 CD4^+^ T cell line. CANX-silencing induced a marked inhibition of CD4^+^ T cell activation by pp65-transfected cells (Fig 7F, left and middle panels). In contrast, the inhibition of CANX expression did not influence the capacity of peptide-loaded cells to activate pp65-specific CD4^+^ T cells (Fig 7F right panel). We show here that T6BP silencing affects CANX functions resulting in CD74 degradation and aberrant MHC-II peptide loading and antigen presentation to CD4^+^ T cells.

## Discussion

We demonstrate here that T6BP regulates the loading and presentation of endogenous viral antigens by MHC-II molecules. This function of T6BP in antigen presentation has a direct implication in the activation of virus specific CD4^+^ T cells. The action of T6BP is broader than what we initially anticipated as it affects the presentation of various antigens whose processing is dependent or independent on autophagy degradation. We further show that T6BP shapes the immunopeptidome of MHC-II molecules. We provide evidence that T6BP controls MHC-II molecule peptide loading in particular through its interaction with the cytosolic tail of CANX that stabilizes the invariant chain. However, this is probably not the only way by which T6BP affects MHC-II restricted antigen presentation as it also participates in the regulation of the trafficking of MHC-II-loading compartments and more globally of acidified vesicular compartments.

We previously reported that HIV-infected cells present MHC-II-restricted HIV Gag- or Env-derived antigens to HIV-specific CD4^+^ T cells (Coulon *et al*., 2016). The processing of these native antigens does not rely on the autophagy pathway. However, when targeted to autophagosomes, using LC3, HIV Gag processing is dependent on autophagosomal degradation. In fact, using DC and HeLa-CIITA cells we showed that as compared to native HIV Gag protein, the targeting of HIV Gag to autophagosomes leads to a more robust activation of Gag-specific T cells (Coulon *et al*., 2016). We concluded that depending on the cellular localisation, the same antigens may be degraded by various endogenous routes leading to MHC-II loading. We reveal here that T6BP modulates Gag antigen presentation independently of its cytosolic or autophagosomal cellular localisation. Likewise, T6BP affects the presentation of an HCMV pp65-derived peptide. Our analysis of the immunopeptidome of HeLa-CIITA cells uncovers that T6BP has a broad influence on the repertoire and relative abundance of the peptides presented by MHC-II molecules. Without excluding the possibility, these observations do not indicate whether T6BP also affects the exogenous pathway of antigen presentation by MHC-II molecules. Indeed, others and we (Fig S2) have demonstrated that the landscape of peptide naturally presented by MHC-II molecules contains a large fraction of peptides derived from intracellular proteins (Muntasell *et al*, 2002; Rammensee *et al*., 1999; Rudensky *et al*., 1991). In our model system, HeLa-CIITA cells, we could not ask whether the silencing of T6BP affects the exogenous pathway because these cells lack the ability to present exogenous viral antigens to CD4^+^ T cells (Coulon *et al*., 2016). This is a weakness but also a strength of this model system since it allowed focusing our work on the endogenous pathway. Nevertheless, we intended to ask whether T6BP might affect antigen presentation by primary APCs. We used several means to silence T6BP in MDDCs including siRNA and shRNA. Unfortunately, whatever the protocol and independently of T6BP expression, the tools used to transfect or to transduce MDDC induced a maturation of the cells and affected the traffic of MHC-II molecules which prohibited from drawing any conclusion on the potential role of T6BP in MDDCs. Nonetheless, our results strongly suggest that at least in epithelial cells that can turn into APCs upon inflammation (Wijdeven *et al*., 2018), T6BP expression shapes the peptide repertoire, the relative abundance and the relative affinity of epitopes presented by MHC-II molecules. Indeed, our immunopeptidomic data suggest that the expression of T6BP favours the loading of peptides with a higher binding capacity to HLA-II molecules. As expected, based on previous work with cell lines (Alvaro-Benito *et al*., 2018) or analysing tissue samples (Marcu *et al*., 2021), we observed a natural variation of the MHC-II immunopeptidome between two mock-treated samples with about 60% of shared peptides. However, in these mock samples, we did not notice a significant difference in terms of relative quantity of core epitopes. This is in sharp contrast to the comparison of wild-type and T6BP-silenced cells where half of the MHC-II binding core epitopes were differentially presented. Remarkably, we also observed a natural variation of the immunopeptidome of MHC-I molecules, with about 50% of shared peptides. Notably, T6BP silencing did not have a strong influence on the repertoire of peptides presented by MHC-I molecules, as the percentage of shared peptides with the control siRNA-treated cells was also around 50%. In addition, the affinity of MHC-I ligands was comparable in wild type cells and cells silenced for T6BP expression. Our results strongly suggest that the action of T6BP is restricted to the MHC-II antigen presentation pathway and affects both the abundance and the affinity of core epitopes to HLA molecules.

The chaperones HLA-DM and the invariant chain (Ii) CD74 tightly regulate the loading of peptides on MHC-II molecules. Previous studies have shown that the levels of expression of both HLA-DM and CD74 affect the repertoire and the affinity of the peptides presented by various HLA-DR and HLA-DQ alleles (Alvaro-Benito *et al*., 2018; Muntasell *et al*., 2004; Ramachandra *et al*, 1996). Remarkably, in the absence of T6BP, we noticed a reduced expression level of CD74 whereas HLA-DM and HLA-DR expression levels remained unchanged. Overall, we did not observe a significant influence of T6BP silencing on the expression levels of other intracellular markers/proteins than CD74. Our analysis of the expression kinetics of MHC-II complexes, using pulse/chase experiments, further showed that the lack of T6BP induces a rapid and exacerbated degradation of CD74. The expression levels of CD74 could be rescued by inhibiting lysosomal acidification but not proteasomal functions suggesting that CD74 is aberrantly degraded in endo-lysosomes in T6BP-depleted cells. These observations probably explain the modifications of the immunopeptidome and the instability of peptide-loaded MHC-II αβ heterodimers, observed in the absence of T6BP expression. However, the diversity of the immunopeptidome could also be due to variations in the origin of the peptides presented by MHC-II molecules (Muntasell *et al*., 2004); in particular since we observed a global modification of the cellular distribution of acidified vesicular compartments in the absence of T6BP. Although we observed, using functional enrichment analysis (Funrich, Fig S2), a slight influence of T6BP silencing on the cellular distribution of the source protein of MHC-II ligands, this cannot by itself explain the effects of T6BP on the peptide repertoire. In particular since, T6BP expression also dictates the relative affinity of pepides presented by MHC-II molecules. Overall, our observations suggest that T6BP expression influences the degradation kinetics of CD74 with a direct influence on the diversity and affinity of the immunopeptidome of MHC-II molecules.

The ER chaperone CANX that participates in the quality control of protein folding in the ER has been shown to play an important role in the assembly of the nonameric MHCαβ-CD74 complexes (Arunachalam & Cresswell, 1995). CANX retains and stabilizes CD74 in the ER until it assembles with the MHC α- and β-chains (Romagnoli & Germain, 1995). It also binds newly synthetized α- and β-chains of HLA molecules until it forms, with CD74, a complete nonamer (Anderson & Cresswell, 1994). Interestingly, it has been shown that the treatment with Tunicamycin or the expression of CD74 mutants, lacking N-linked glycosylation, both impeding the interactions with CANX, can induce CD74 degradation (Romagnoli & Germain, 1995). We corroborate here that CANX influences CD74 expression. Using siRNA silencing, we provide a direct demonstration that the lack of CANX expression induces CD74 degradation. Furthermore, we show that the silencing of CANX in APC diminishes their capacity to activate antigen-specific CD4^+^ T cells. Remarkably, using co-immunoprecipitation followed by mass spectrometry, we identified CANX as a binding partner of T6BP, among 116 high-confidence T6BP proximal proteins. T6BP and CANX interaction was confirmed in HeLa-CIITA as well as in B cells and DC, strongly suggesting that our observations are not limited to our model APCs.

It has been proposed that the ER distribution of CANX might determine its functions as chaperone involved in protein quality control or in the regulation of Ca2+ transfer to mitochondria (Lynes *et al*, 2013). Several post-translational modifications influence both the cellular distribution and the functions of calnexin: phosphorylation of the cytosolic tail of CANX leads to a re-distribution from peripheral ER tubules to a juxtanuclear ER (Myhill *et al*, 2008) while palmitoylation of the tail seems to assign specific tasks to calnexin within the ER in particular a role in protein folding (Lynes *et al*., 2013) and a localization at the proximity of the ribosome complexes (Lakkaraju *et al*, 2012). We show here that T6BP binds to the cytosolic tail of CANX. To our knowledge, there is no published evidence that CANX is ubiquitinated on its cytosolic tail but we believe this possibility should be investigated further as T6BP might use its Ub-binding domain to bind to CANX, providing a molecular link between T6BP and CANX functions in the formation of MHC-II loading complexes and thus antigen presentation. Note that CANX has also been shown to contribute to the assembly of MHC-I molecules by protecting free MHC-I molecules from degradation (Vassilakos *et al*, 1996) and by binding to the TAP-tapasin complex (Diedrich *et al*, 2001). However, CANX action on MHC-I molecule assembly and transport to the cell surface seems dispensable (Prasad *et al*, 1998; Scott & Dawson, 1995) which would explain the lack of influence of CANX on the MHC-I immunopeptidome, in the absence of T6BP.

We suggest here that CANX is a key factor involved in T6BP-mediated regulation of MHC-II loading that strongly influences the repertoire and affinity of presented peptides. However, we do not exclude that other cellular factors or pathways also participate in T6BP action on MHC-II molecules. Jongsma et al showed that the ER ubiquitin ligase RNF26 interacts with p62 (SQSTM1) and that it employs a ubiquitin-based communication with Tollip, T6BP, or EPS15 to cluster recycling, early and late endosomes in so-called perinuclear clouds. In the model, deubiquitination of p62 by the DUB UPS15 leads to the release of the vesicles for maturation (Jongsma *et al*., 2016). The authors showed that the silencing of RNF26 leads to the dispersion of EEA1+ and CD63+ vesicles, in the cytoplasm of the cells. They also show that the silencing of T6BP has a broad influence on the shape of cells and leads to the dispersion of the cloud of CD63+ vesicles. Our observations also highlight that T6BP-silencing influences CD63+ vesicles, however our results indicate that CD63+ vesicles are maintained in the perinuclear region. We confirmed these observations using lysotracker and LAMP-1 markers of acidified and late endosomes respectively. The different behaviour of CD63+ vesicles in siT6BP-treated cells might be due to the expression of the HLA locus driven by CIITA, in our model of APCs. Nonetheless, our work confirms the involvement of T6BP in the traffic of endosomes. In addition, we found, in the interactome study of T6BP, the DUB UPS15. Interestingly, although T6BP affects EEA1+ early endosomes perinuclear localization, we excluded the possibility that the effect of T6BP on MHC-II-restricted antigen presentation relies on early endosomal defect since EEA1+ vesicles do not contain MHC-II molecules in our APCs. Overall, we show that T6BP impacts the trafficking of MHC-II-rich compartments that bear hallmarks of the MIIC, namely HLA-DM, CD63+ and LAMP-1. More globally T6BP-silencing seems to affect the cellular localization of acidified vesicular compartments corresponding to late endosome, lysosomes and as already demonstrated autophagosomes (Petkova *et al*., 2017), without significantly reducing the expression levels of the various markers. Accumulating evidences suggest that the positioning of intracellular vesicles controls their functions (Neefjes *et al*, 2017) and intravesicular pH in particular (Johnson *et al*, 2016). The repositioning of the MIIC mediated by T6BP could influence the activation of intravesicular pH-dependent proteases that participate in the cleavage of CD74 and antigen processing. To this regard, it is interesting to note that we identified in the interactome of T6BP, ATP6V0A1 a subunit of the vATPase that plays a critical role in mediating vesicular acidification (Forgac, 2007). On the other hand, CD74 has been shown to play, by itself, a role in endosomal membrane trafficking (Margiotta *et al*, 2020; Romagnoli *et al*, 1993) and perhaps in proteolysis of antigens (Schröder, 2016). It was recently reported that CD74 p41 isoform interacts with cathepsins and regulates their activity in endosomes (Bruchez *et al*, 2020). These studies and others suggest that CD74 might exert various functions not limited to its canonical role in antigen presentation and as a chaperone (Schröder, 2016). Thereafter, the modifications in late endosomes/lysosomes positioning and of the MHC-II immunopeptidome in T6BP-silenced cells could be an indirect consequence of CD74 degradation.

Our interactome also revealed as candidate partners for T6BP, the HLA-A, -C, -DQα1 and -DRβ1 molecules. Interestingly, the cellular distribution of both HLA class-I and -II molecules is regulated by the addition of ubiquitin (Ub) moieties (De Angelis Rigotti *et al*, 2017; Shin *et al*, 2006; Walseng *et al*, 2010). In the case of HLA-DR, ubiquitination of the lysine residue (K235) of HLA-DRβ1 plays a major role in regulating the cellular localisation of mature HLA-DR molecules (Lapaque *et al*, 2009). Although T6BP has the capacity to bind ubiquitinated proteins, we favour the hypothesis that T6BP and HLA molecules might be part of larger molecular complexes or lipid compartments in particular since we observed neither a co-localization of T6BP with HLA-DR molecules nor a significant difference in the kinetics of HLA-DR endocytosis in HeLa-CIITA cells silenced for T6BP expression.

The autophagy receptor T6BP as well as OPTN, NDP52, p62 and NBR1 harbour Ub- and LC3-binding motifs, whose functions are well characterized in selective autophagy (Kirkin & Rogov, 2019). In addition, T6BP and NDP52 share a N-terminal SKIP carboxyl homology (SKICH) domain target of TANK-binding kinase-1 (TBK1) phosphorylation that acts as an upstream regulator of mitophagy (Fu *et al*, 2018). Beyond selective autophagy, ARs also modulate the traffic and maturation of autophagosomes and endosomes. The Ub-binding domains of T6BP, OPTN and p62 have been implicated in regulating the NF-κB pathway where together with A20 they down-modulate the activation of this pathway during inflammation (Weil *et al*, 2018). T6BP, NDP52 and OPTN also bind to Myosin-VI, which recruits Tom-1-expressing endosomes and lysosomes, facilitating autophagosome maturation in lytic vesicles (Morriswood *et al*., 2007; Sahlender *et al*., 2005; Tumbarello *et al*., 2012). A recent work suggested that Ub and Myosin-VI compete for the binding to T6BP (Hu *et al*, 2018). In T6BP’s interactome, we identified known protein partners of T6BP involved for instance in NF-κB signalling pathways (e.g. TBK1, TRAF2, ITCH, SQSTM1/p62). However, other described T6BP partners such as Myosin-VI were not identified, likely reflecting that T6BP interactions occur in a cell type-specific manner (O’Loughlin *et al*, 2018). Nonetheless, we revealed novel potential partners of T6BP involved in the Ubiquitin/Proteasome (e.g. E3 Ub-ligases UBR1, 2 and 4), the UPR/protein folding (e.g. BAG6, EIF2a, calnexin), endocytosis (e.g. COPB1) and antigen presentation pathways (e.g. HLA molecules). We also found that T6BP interacts with NBR1 that has been found together with p62 in larger protein complexes and ubiquitinated protein aggregates (Weil *et al*., 2018). Much remains to be learned on the molecular mechanisms and motifs involved in the variety of T6BP molecular interactions and functions, in particular in MHC-II antigen presentation that we unravel here. Side by side comparison of AR functional domains will most likely bring new insights.

Our work reveals that T6BP regulates the cellular positioning of the MHC-II loading compartment and the stability of CD74, through an interaction with CANX, and exert a direct influence on MHC-II-restricted antigen presentation. This novel role of T6BP in the activation of adaptive antiviral immunity further highlights the diverse non-redundant functions exerted by autophagy receptors.

## Materials and Methods

### Cells

HeLa-CIITA cells were provided by Philippe Benaroch (Institut Curie, Paris, France), are homozygotes for HLA-DRβ1*0102 allele, and were cultured with RPMI GlutaMax 1640 (Gibco) complemented with 10% FBS (Dutscher), 1% Penicillin/Streptomycin, 50µg/mL Hygromycin B (Thermo Fisher).

### HIV-1-& HCMV-specific CD4^+^ T cell clones

Gag-specific CD4^+^ T cell clones (F12) are specific for HIV Gag-p24 (gag2: aa 271-290) and restricted by HLA-DRβ1*01 as previously described (Coulon *et al*., 2016; Moris *et al*., 2006). HCMV-specific CD4^+^ T cell clones are specific for HCMV pp65 antigen (pp65: aa 108-127) and restricted by HLA-DRβ1*01. Pp65-specific clones were isolated from PBMCs of healthy donors after several round of *in-vitro* stimulation with synthetic peptide corresponding to immunodominant epitopes from the pp65 protein. Pp65-specific cells were isolated using the IFN-γ secreting assay from Miltenyi Biotec and cloned by limiting dilution. F12 and pp65 clones were restimulated and expanded, as previously described (Moris *et al*., 2006), using irradiated feeders and autologous or HLA-matched lymphoblastoid cell lines loaded with cognate peptides in T cell cloning medium: RPMI 1640 containing 5% human AB serum (Institut Jacques Boy), recombinant human IL-2 (100 IU/ml, Miltenyi Biotec), PHA (0,25 µg/ml, Remel), non-essential amino acids, and sodium pyruvate (both from Life Technologies). At least 1h before coculture with HeLa-CIITA cells, T cell clones were thawed and allowed to rest at 37°C in RPMI containing DNAse (5µg/mL, New England Biolabs).

### Viral antigens and plasmids

The pTRIP-CMV-Gag (a kind a gift from Nicolas Manel (Institut Curie, Paris, France), pGag-LC3 and Gag-LC3_G120A_ plasmids were already described (Coulon *et al*., 2016). The pp65 encoding cDNA (a kind gift from Xavier Saulquin, Université de Nantes, Nantes) was cloned in the lentiviral vector cppT-EF1α-IRES-GFP. The GFP-T6BP encoding plasmid is a kind gift from Folma Buss (University of Cambridge, UK) (Tumbarello *et al*., 2015).

### Cell transfections

HeLa-CIITA cells were incubated in 6-well plates using 2-4.10^5^ cells/well using OPTIMEM (Gibco) complemented with 10% FBS, 1% Penicillin/Streptomycin. Twenty-four hours later, cells were transfected with 40 pmol of siRNA targeting NDP52 (L-010637-00-0005), OPTN (L-016269-00-0005), p62 (L-010230-00-0005), T6BP (L-016892-00-0020 Dharmacon or SI02781296, Qiagen), CANX (SI02663367 and SI02757300, Qiagen) or a scrambled siRNA as control (D-001810-10-20, Dharmacon), using Lipofectamine RNAiMax (13778-150, Thermo Fisher) as transfection reagent. After 24h of transfection, cells were transfected with the cDNA encoding the viral antigens (1µg per well of a 6-well plate) using Viromer RED (Lipocalyx) and following manufacturer instructions. Twenty-four hours later, Gag and pp65 expressions were assessed using anti-Gag antibody (KC57-RD1, Beckman-Coulter) and anti-pp65 antibody (mouse, Argene) combined with goat anti-mouse antibody (AF488, Thermo Fisher), respectively.

### Flow cytometry

Cell viability was evaluated using LIVE/DEAD (Thermo Fisher) or Zombie (Biolegend) and the following antibodies were used: HLA-DR specific L243 and TÜ36 (both in house and kindly provided by Philippe Benaroch, Institut Curie, Paris) and goat anti-mouse (AF488, Thermo Fisher). Forty-eight hours after siRNA transfection, cell-surface staining assays were performed using standard procedures (30min, 4°C). HIVGag and HCMVpp65 production was detected using intracellular staining. Briefly, cells were fixed with 4% PFA (10min, RT), washed, and permeabilized with PBS containing 0,5% BSA and 0,05% Saponin, prior to antibody staining. Samples were processed on Fortessa cytometer using FACSDiva software (BD Biosciences) and further analysed using FlowJo2 software (Tree Star).

### IFN-γ ELISPOT assay

ELISPOT plates (MSIPN4550, Millipore) were pre-wet and washed with PBS, and coated overnight at 4°C with anti-IFN-γ antibody (1-DIK, Mabtech). Plates were washed using PBS and then saturated with RPMI complemented with 10% FBS. Plates were washed and HeLa-CIITA cells (10^5^ cells/well) were co-cultured with T cell clones (5.10^3^ and 1.10^3^ cells/well) overnight at 37°C. Cells were removed and plates were then washed with PBS-0,05% Tween-20 prior incubation with biotinylated anti-IFN-γ antibody (7-B6-1, Mabtech) (2h, RT). Spots were revealed using alkaline-phosphatase coupled to streptavidin (0,5U/ml, Roche Diagnostics) (1h, RT) and BCIP/NBT substrate (B1911, Sigma-Aldrich) (30min, RT). Reactions were stopped using water. Number of spots were counted using AID reader (Autoimmun Diagnostika GmbH). For each experimental condition, ELISPOTs were performed mostly in triplicates or at least in duplicates.

### Western Blotting

Forty-eight hours after siRNA transfection, 10^6^ HeLa-CIITA cells were washed in cold PBS and lysed in 100-300µl of lysis buffer (30min, 4°C), mixing every 10 min. When stated, cells were pre-treated with Chhloroquine (50µM, 32h) or Epoxomicin (50nM, 32h). Depending on the experiments, two different lysis buffers were used: 1-PBS containing 1% Nonidet P-40; 2-50 mM Tris-HCl pH 7.5 containing 100 mM NaCl, 1% Triton X-100, 0.5 mM EGTA, 5 mM MgCl, 2 mM ATP, both supplemented with 1x Protease Inhibitor (Roche). Cell lysates were then centrifuged at 20,000g (10 min, 4°C), supernatants harvested and mixed with Sample Buffer (NuPAGE, Invitrogen) and Sample Reducing Agent (NuPAGE, Invitrogen) and denatured (5min, 95°C). Denatured samples were analysed by SDS gel electrophoresis using 4-12% Bis-Tris gels (NuPAGE, Invitrogen), transferred to a nitrocellulose membrane (NuPAGE, Invitrogen) and immunoblotted. Anti-T6BP (HPA024432, Sigma-Aldrich), anti-NDP52 (HPA023195, Sigma-Aldrich), anti-CANX (PA5-34754, ThermoFisher), anti-OPTN (ab23666, Abcam), anti-p62 (sc-28359, Santa Cruz Biotechnology), anti-HLA-DR (TAL1B5, Invitrogen), anti-LC3 (M152, MBL International), anti-actin (3700S, Cell Signaling Technology), anti-tubulin (2148S, Cell Signaling Technology), anti-CD74 (ab22603, Abcam), anti-GFP (11814460001, Roche), goat anti-mouse (Sigma-Aldrich), and goat anti-rabbit both coupled to HRP (Abcam) antibodies were used according to manufacturer instructions. Blots were revealed using Pierce ECL Plus Substrate (Invitrogen) and chemiluminescence analysed using ImageQuant LAS 4000.

### Confocal microscopy

Forty-eight hours after siRNA transfections, HeLa-CIITA cells were plated on glass coverslips and then fixed with 4% PFA (10min, RT). Cells were washed 3 times with PBS, saturated with goat or donkey serum and permeabilized with PBS containing 0,5% BSA and 0,05% Saponin (1h, RT). Cells were washed with PBS and incubated (OVN, 4°C) with primary antibodies: L243 or TÜ36 (both in house and kindly provided by P. Benaroch, Institut Curie, Paris), rabbit anti-HLA-DR (a kind gift from Jack Neefjes), LAMP-1 (H4A3, DSHB), CD63 (MA-18149), anti-CD74 (14-0747-82) all from ThermoFisher, anti-EEA1 (C45B10, Cell signalling), T6BP (HPA024432, Sigma-Aldrich) and Lysotracker (Deep Red, L12492). Cells were incubated with species specific antibodies: goat anti-mouse coupled to Alexa Fluor 488 or Alexa Fluor 405 (ThermoFischer), donkey anti-rabbit coupled to Alexa 546 (A10040, Invitrogen) in PBS containing 0,5% BSA and 0,05% Saponin (1h, RT). When required sequential stainings were performed. Nuclei were stained with DAPI (17507, AAT Bioquest). After washing with PBS, samples were mounted on glass slides with Dako fluorescence mounting medium. Samples were imaged using a laser scanning confocal microscope with 63X, NA 1.3 oil immersion objective. The number of vesicles, the intensity and the distances of each vesicle to nucleus were quantified using an in-house ImageJ Python script (developed by Aziz Fouché, ENS Paris-Saclay, Paris). Potential co-localizations were determined using the object based co-localization method JACoP (Just Another Co-localization Plugin) and coloc2 (Pearson’s coefficient) of the ImageJ software, for punctuated/vesicular and cytosolic/diffuse staining, respectively. For the Proximity ligation assay (PLA), 48h after siRNA transfections, HeLa-CIITA cells, cultured overnight on coverslips, were fixed with 4% PFA (10min, RT), washed 3x with PBS and treated with 50mM NH4Cl (15min, RT) prior incubation with anti-T6BP (Rabbit, 1:200, HPA024432) and anti-CANX (Mouse, 1:500, MA3-027, Thermo fisher scientific) in PBS containing 1%, BSA and 0.1% Saponin (1h, RT). PLA secondary probes (DUO92002, DUO92004) were then used according to the manufacturer’s instructions (Sigma-Aldrich). Briefly, 40 µl (1:5 dilution) of the PLA were added (37 °C, 60 min), washed with buffer A (DUO82049, Sigma-Aldrich). 40 µl of the ligation mix (DUO92014, Sigma-Aldrich) was then applied to each of the coverslips to complete the ligation process (30min, 37 °C). Coverslips were then incubated with 40µl polymerisation mix (100min, 37 °C) and washed. Coverslips were mounted on slides with Fluoromount G (Thermo fisher). Images were acquired using SP8 confocal microscope and dots were counted using find maxima plugin in Fiji software.

### Reverse transcription quantitative Polymerase Chain Reaction (RT-qPCR)

Total cellular RNA was isolated using RNeasy kit (Qiagen, Valencia, CA). RNA concentrations were determined by spectrophotometry at 260 nm. The relative level of CD74 and T6BP mRNAs were determined using the comparative Cq method. Actin was used as endogenous control. The primers and probes used for quantitation of CD74, T6BP and actin were designed by Olfert Landt and purchased from TIB MolBiol. Sequences are listed in the table below. The RT-qPCRs were performed in a Light Cycler 1.5 instrument in capillaries using a final volume of 20 µl. The reactions were performed using 300 nM specific sense primer, 300 nM specific antisense primer, 200 nM specific TaqMan probe (TM) and the LightCycler® Multiplex RNAVirus Master mix (ROCHE). The programs were: reverse-transcription 55°C for 10min, initial denaturation 95°C for 5min followed by 45 cycles of amplification. For CD74 and T6BP cycles were: 95°C for 5s, 60°C for 15s + fluorescence measurement and 72°C for 5s. For actin cycles were: 95°C for 20s, 67°C for 30s + fluorescence measurement and 72°C for 5s.

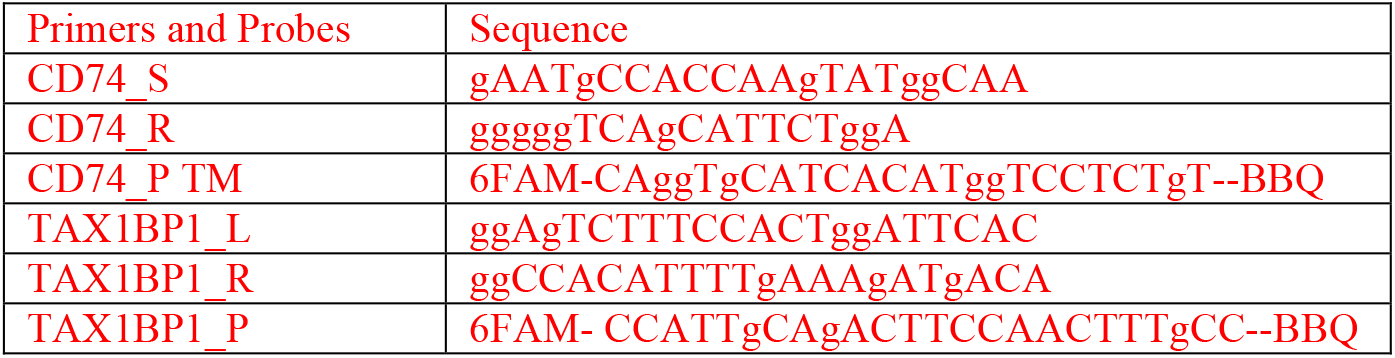
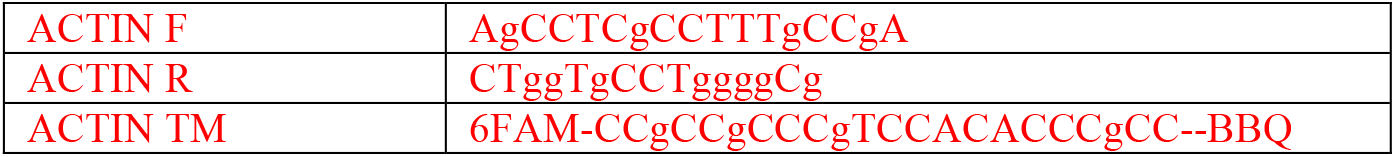

### Pulse-Chase experiment

48h post siRNA transfection HeLa-CIITA cells were preincubated in Met/Cys-free RPMI 1640 medium containing 1% Penicillin/Streptomycin, 1% glutamine and 10% dialyzed FCS for 1h. Cells (five million) were then pulsed for 30 min with 0.5 mCi of ^35^S-Met/Cys (ICN) and chased in unlabelled medium supplemented with 5 mM cold methionine. Cells were collected at the indicated time points and then lysed in a buffer containing 20 mM Tris-HCl (pH 7.5), 150 mM NaCl, 2 mM MgCl2, 1% Triton X-100, and an inhibitor cocktail (Roche). Each sample was normalised for the protein concentration. Lysates were precleared with mouse serum, and MHC class II/Ii complexes were immunoprecipitated with the TÜ36 antibody and/or reimmunoprecipitated with VICY1 antibody (14-0747-82, ThermoFischer). Samples were boiled (95°C) or unboiled (room temperature) in SDS loading buffer and separated on 12% SDS-PAGE (Novex). Quantification of the results was made using a phosphoimager (Fuji).

### Electron Microscopy

HeLa-CIITA cells (2.10^5^ cells/well) were cultured on glass cover slips in 6-well plates. 24h later, cells were transfected as described with siRNA Control or targeting T6BP, cells were cultured for 48 hours and fixed with 2.5% Glutaraldehyde, 1% PFA for 1 h at room temperature. The cover slips were washed 3 times with 0.2 M phosphate buffer pH 7.4, followed by a 1-hour incubation in 1% Osmium, 1.5% Ferrocyanide of Potassium. After 3 washes in water, the cover slips were successively treated with 50%, 70%, 90%, 100% and 100% Ethanol for 10 minutes each. The cover slips were then incubated for 2 hours on 50% epoxy in ethanol, followed by 2 hours in pure epoxy, and finally in pure epoxy overnight for polymerization at 60°C. Ultrathin (70 nm) sections were cut using a diamond knife (45° angle) on a Leica UC6 ultramicrotome. Sections were collected on Formvar™ carbon-coated copper grids. Some sections were stained with uranyl acetate at 2% (Merk) for 15 min and lead Citrate (Agar) and washed three times with milliQ water and dried at room temperature. Observations were performed with a JEOL JEM-1400 transmission electron microscope operating at 120 kV. Images were acquired using a post-column high-resolution (9 10^6^ pixels) camera (Rio9; Gatan) and processed with Digital Micrograph (Gatan) and ImageJ.

### Immunopeptidome

#### Isolation of HLA ligands

HLA class-I and -II molecules of HeLa-CIITA cells were isolated using standard immunoaffinity purification (Falk *et al*, 1991; Nelde *et al*, 2019). Snap-frozen samples were lysed in 10 mM CHAPS/PBS (AppliChem, Gibco) with 1x protease inhibitor (Roche). HLA class-I and -II-associated peptides were isolated using the pan-HLA class I-specific mAb W6/32 and the pan-HLA class II-specific mAb TÜ39 (both in house mouse monoclonal) covalently linked to CNBr-activated Sepharose (GE Healthcare). HLA-peptide complexes were eluted by repeated addition of 0.2% TFA (trifluoroacetic acid, Merck). Eluted HLA ligands were purified by ultrafiltration using centrifugal filter units (Millipore). Peptides were desalted using ZipTip C18 pipette tips (Millipore), eluted in 35 µl 80% acetonitrile (Merck)/0.2% TFA, vacuum-centrifuged and resuspended in 25 µl of 1% acetonitrile/0.05% TFA and samples stored at -20 °C until LC-MS/MS analysis.

#### Analysis of HLA ligands by LC-MS/MS

Isolated peptides were separated by reversed-phase liquid chromatography (nano-UHPLC, UltiMate 3000 RSLCnano; ThermFisher) and analysed in an online-coupled Orbitrap Fusion Lumos mass spectrometer (Thermo Fisher). Samples were analysed in five technical replicates and sample shares of 20% trapped on a 75 µm × 2 cm trapping column (Acclaim PepMap RSLC; Thermo Fisher) at 4 µl/min for 5.75 min. Peptide separation was performed at 50 °C and a flow rate of 175 nl/min on a 50 µm × 25 cm separation column (Acclaim PepMap RSLC; Thermo Fisher) applying a gradient ranging from 2.4 to 32.0% of acetonitrile over the course of 90 min. Samples were analysed on the Orbitrap Fusion Lumos implementing a top-speed CID method with survey scans at 120k resolution and fragment detection in the Orbitrap (OTMS2) at 60k resolution. The mass range was limited to 400–650 m/z with precursors of charge states 2+ and 3+ eligible for fragmentation.

#### Database search and spectral annotation

LC-MS/MS results were processed using Proteome Discoverer (v.1.3; Thermo Fisher) to perform database search using the Sequest search engine (Thermo Fisher) and the human proteome as reference database annotated by the UniProtKB/Swiss-Prot. The search-combined data of five technical replicates was not restricted by enzymatic specificity, and oxidation of methionine residues was allowed as dynamic modification. Precursor mass tolerance was set to 5 ppm, and fragment mass tolerance to 0.02 Da. False discovery rate (FDR) was estimated using the Percolator node (Käll *et al*, 2007) and was limited to 5%. For HLA class-I ligands, peptide lengths were limited to 8–12 amino acids. For HLA class-II, peptides were limited to 12–25 amino acids of length. HLA class-I annotation was performed using NetMHCpan 4.0 (Jurtz *et al*., 2017) annotating peptides with percentile rank below 2% as previously described (Ghosh *et al*, 2019).

For HLA class-II peptides, the Peptide Lansdscape Antigenic Epitope Alignment Utility (PLAtEAU) algorithm (Alvaro-Benito *et al*., 2018) was used to identify and to estimate the relative abundance of the core epitopes based on the LC-MS/MS intensities. The results are presented as Volcano plots using Perseus software (Tyanova *et al*, 2016). The relative affinities of the core epitope to HLA-DRβ1*0102, expressed by HeLa-CIITA cells, was estimated using NetMHCIIpan 4.0 (Reynisson *et al*., 2020).

### Interactome

#### Co-immunoprecipitation

HeLa-CIITA cells were harvested 24h after cDNA transfection with either plasmid encoding GFP-T6BP (kind gift from F.Buss, Cambridge, UK) or encoding GFP. 2.10^7^ cells were washed in cold PBS and lysed in 300 µl of lysis buffer (50 mM Tris-HCl pH 7.5, 100 mM NaCl, 1% Triton X-100, 0.5 mM EGTA, 5 mM MgCl, 2 mM ATP, and 1x Protease Inhibitor (Roche) (30min, ice), mixing every 10min and centrifuged at 20,000g (20min, 4°C). Lysates were recovered, 300µl of wash buffer (50 mM Tris-HCl pH 7.5, 150 mM NaCl, 0.5 mM EDTA) was added and the pellets were discarded. GFP-Trap agarose magnetic beads (Chromotek) were vortexed, 25µl of bead slurry was washed 3 times with cold wash buffer. Each diluted lysate was added to 25 µl of equilibrated beads and tumbled end-over-end (1h, 4°C). Beads were collected using a magnetic support and washed 3 times. For Western Blot analysis, SDS-sample buffer was added to aliquots and the samples were boiled (5 min, 99°C).

#### On-bead digestion for mass spectrometry

Following immunoprecipitation with GFP-Trap (Chromotek), digestions were performed using manufacturer instructions on the P3S proteomic core facility of Sorbonne Université.

For each sample, beads were resuspended in 25 µl of elution buffer I (50 mM Tris-HCL pH 7.5, 2M urea, 5 µg/ml sequencing grade Trypsine, 1 mM DTT) and incubated in a thermomixer at 400rpm (30min, 30°C). Beads were collected using a magnetic support and the supernatants were recovered. For elution, beads were then washed with 50 µl of elution buffer II (50 mM Tris-HCL pH 7.5, 2 M urea, 5 mM iodoacetamide) and collected with a magnetic support. Supernatants were harvested and mixed with the previous ones. This elution was repeated once. Combined supernatants were incubated in a thermomixer at 400 rpm (overnight, 32°C). Reactions were stopped by adding 1µl trifluoroacetic acid and digests were desalted using home-made StageTips. StageTips were first rehydrated with 100µl of methanol and then equilibrated with 100 µl of 50% acetonitrile 0.5% acetic acid. After peptide loading, StageTips were washed with 200 µl of 0.5% acetic acid and peptides were eluted with 60 µl of 80% acetonitrile 0.5% acetid acid. Eluted peptides were totally dried using a SpeedVac vacuum concentrator (Thermo), solubilised in 20 µl of 2% acetonitrile 0.1% formic acid before LC-MS/MS analysis.

#### LC-MS/MS

Peptide mixtures were analysed with a nanoElute UHPLC (Bruker) coupled to a timsTOF Pro mass spectrometer (Bruker). Peptides were separated on an Aurora RP-C18 analytical column (25 cm, 75 µm i.d., 120 Å, 1,6 µm IonOpticks) at a flow rate of 300 nL/min, at 40°C, with mobile phase A (ACN 2% / FA 0.1%) and B (ACN 99.9% / FA 0.1%). A 30 min elution gradient was run from 0% to 3% B in 1 min, 3% to 15 % B in 17 min then 15% to 23% B in 7 min and 23% to 32% B in 5 min. MS acquisition was run in DDA mode with PASEF. Accumulation time was set to 100 msec in the TIMS tunnel. Capillary voltage was set to 1,6 kV, mass range from 100 to 1700 m/z in MS and MS/MS. Dynamic exclusion was activated for ions within 0.015 m/z and 0.015 V.s/cm^2^ and released after 0,4 min. Exclusion was reconsidered if precursor ion intensity was 4 times superior. Low abundance precursors below the target value of 20,000 a.u and intensity of 2,500 a.u. were selected several times for PASEF-MS/MS until the target value was reached. Parent ion selection was achieved by using a two-dimensional m/z and 1/k0 selection area filter allowing the exclusion of singly charged ions. Total cycle time was 1,29 sec with 10 PASEF cycles.

#### Data Analysis

Raw data were processed with MaxQuant version 1.6.5.0, with no normalisation, no matching between runs and with a minimum of 2 peptide ratios for protein quantification. The output protein file was filtered with ProStar 1.14 to keep only proteins detected in 2 samples or more in at least 1 of the 2 conditions. Missing values were imputed using SLSA (Structured Least Square Adaptative) algorithm for partially missing values in each condition and DetQuantile algorithm for missing values in an entire condition. In order to select relevant binding partners, data were statistically processed using limma test and filtered to retain only differentially expressed preys (FDR 1%) with a fold change ≥ 10 between T6BP-GFP and GFP conditions. Selected preys were uploaded to the CRAPome v2 (Contaminant Repository for Affinity Purification) online analysis tool to identify potential contaminants. For each binding partners a Significance Analysis of INTeractome (SAINT) probability threshold was assessed by the Resource for Evaluation of Protein Interaction Networks (REPRINT) using the default settings. Selected preys were then uploaded in Ingenuity Pathway Analysis software version 49932394 (QIAGEN Inc., https://www.qiagenbioinformatics.com/products/ingenuity-pathway-analysis) to perform annotation and over-representation analysis. Finally, network visualization was designed using Cytoscape software (v. 3.7.1).

**Co-immunoprecipitation:** CANX / T6BP / GST-TM-cytoTail

HeLa-CIITA cells were harvested, counted and lysed in m-RIPA buffer (1% NP40, 1% sodium deoxycholate, 150mM NaCl, 50mM Tris pH 7.5, Complete protease inhibitors; Roche, Basel, Switzerland). CANX and T6BP were immunoprecipitated using anti-CANX antibody PA5-34754 (Invitrogen) coupled to protein A Sepharose magnetic beads (Ademtech 0423) and anti-TAX1BP1 antibody HPA024432 (Sigma-Aldrich) together with protein A Sepharose (GE Healthcare), respectively. The GST-TM-cytoTail nucleotide sequence was synthetized and cloned into pLV-EF1a-IRES-Puro by GeneCust. pLV-EF1a-IRES-Puro was a gift from Tobias Meyer (Addgene plasmid # 85132; http://n2t.net/addgene:85132; RRID:Addgene_85132) (Hayer *et al*, 2016). GST-TM-cytoTail sequence contains the following sequence: CANX leader, GST, a short fragment of the ER luminal domain of CANX, CANX TM, CANX cytosolic tail; and is: ATGGAAGGGAAGTGGTTGCTGTGTATGTTACTGGTGCTTGGAACTGCTATTGTTGAGGCTTCCCCTATACTAGGTTATTGGAAAATTAAGGGCCTTGTGCAACCCACTCGACTTCTTTTGGAATATCTTGAAGAAAAATATGAAGAGCATTTGTATGAGCGCGATGAAGGTGATAAATGGCGAAACAAAAAGTTTGAATTGGGTTTGGAGTTTCCCAATCTTCCTTATTATATTGATGGTGATGTTAAATTAACACAGTCTATGGCCATCATACGTTATATAGCTGACAAGCACAACATGTTGGGTGGTTGTCCAAAAGAGCGTGCAGAGATTTCAATGCTTGAAGGAGCGGTTTTGGATATTAGATACGGTGTTTCGAGAATTGCATATAGTAAAGACTTTGAAACTCTCAAAGTTGATTTTCTTAGCAAGCTACCTGAAATGCTGAAAATGTTCGAAGATCGTTTATGTCATAAAACATATTTAAATGGTGATCATGTAACCCATCCTGACTTCATGTTGTATGACGCTCTTGATGTTGTTTTATACATGGACCCAATGTGCCTGGATGCGTTCCCAAAATTAGTTTGTTTTAAAAAACGTATTGAAGCTATCCCACAAATTGATAAGTACTTGAAATCCAGCAAGTATATAGCATGGCCTTTGCAGGGCTGGCAAGCCACGTTTGGTGGTGGCGACCATCCTCCAAAAAAGAAAGCTGCTGATGGGGCTGCTGAGCCAGGCGTTGTGGGGCAGATGATCGAGGCAGCTGAAGAGCGCCCGAAGAAAGCTGCTGATGGGGCTGCTGAGCCAGGCGTTGTGGGGCAGATGATCGAGGCAGCTGAAGAGCGCCCGTGGCTGTGGGTAGTCTATATTCTAACTGTAGCCCTTCCTGTGTTCCTGGTTATCCTCTTCTGCTGTTCTGGAAAGAAACAGACCAGTGGTATGGAGTATAAGAAAACTGATGCACCTCAACCGGATGTGAAGGAAGAGGAAGAAGAGAAGGAAGAGGAAAAGGACAAGGGAGATGAGGAGGAGGAAGGAGAAGAGAAACTTGAAGAGAAACAGAAAAGTGATGCTGAAGAAGATGGTGGCACTGTCAGTCAAGAGGAGGAAGACAGAAAACCTAAAGCAGAGGAGGATGAAATTTTGAACAGATCACCAAGAAACAGAAAGCCACGAAGAGAGTGA. Hela-CIITA cells were transfected with GST-TM-cytoTail construct. 24h post-transfection, cells were harvested and lysed using NP40 lysis buffer (10mM Tris HCl pH 7.5, 150 mM NaCl, 0,5 mM EDTA, 0,5% IGEPAL). The GST-immunoprecipitation was performed using GST-Trap Agarose from Chromotek. The immunoprecipitation (IP) fractions were extensively washed (at least 3 times) in NP40 lysis buffer. The input, the flow through and the IP fractions were analysed by SDS-PAGE and the proteins revealed using Western blot.

### Statistical analysis

Statistical significances (p-values) were calculated using Prism Software (GraphPad).

## Data Availability

The mass spectrometry proteomics data have been deposited to the ProteomeXchange Consortium via the PRIDE (Perez-Riverol *et al*, 2019) partner repository with the dataset identifier - PXD024330 and 10.6019/PXD024330 - and PXD024417 for the T6BP-interactome and Hela-CIITA immunopeptidome, respectively.

Reviewer account details:

Username

Password: aCQuFy8B

## Acknowledgements

This work was granted by the ANR project AutoVirim (ANR-14-CE14-0022). We thank the “P3S” core facility of Sorbonne University for their expertise. The present work has benefited from the Imagerie facility of Imagerie-Gif, (http://www.i2bc.paris-saclay.fr), member of IBISA (http://www.ibisa.net), supported by l’Agence Nationale de la Recherche (ANR-11-EQPX-0029/Morphoscope), “France-BioImaging” (ANR-10-INSB-04-01) and the Labex “Saclay Plant Science” (ANR-11-IDEX-0003-02). C.R. and M.P. were supported by AutoVirim. We thank the Dormeur Foundation, Vaduz, for providing the AID ELISPOT Reader. We also thank the Agence Nationale de Recherche sur le SIDA et les hepatites virales (ANRS) and Sidaction for fundings. M.P. and L.B. were supported by Sidaction. A.K. and R.J.-M. were ANRS fellows.

## Author contributions

A.M. conceived and designed the project with contributions from B.M.. Design and performed the experiments: M.P., C.R., G.S., A.K., M.G., L.B., C.P., S.G., F.S., R.J-M., E.R., B.M. and B.C.R.. Analysed the data: M.P., C.R., G.S., A.K., L.B., M.G., M.L., F.S., O.D., C.P., S.G., B.M., S.G.-D., B.C.R. and A.M.. Contributed reagents/materials/analysis tools: M.F., A.E., S.S., O.D. and B.M.. Wrote the paper: M.P., C.R., G.S, B.M., B.C.R. and A.M. with contributions from all authors.

## Conflict of interest

All other authors declare no financial or commercial conflict of interest.

## Notes

### Competing Interest Statement

The authors have declared no competing interest.

### Summary of Updates

In the revised version, we provide mechanistic insights by showing that T6BP controls MHC-II molecule peptide loading in particular through its interaction with the cytosolic tail of Calnexin that stabilizes the invariant chain. We extended our observations to professional antigen presenting cells. and more…

