## Supplementary material for "The Autophagy Receptor TAX1BP1 (T6BP) is a novel player in antigen presentation by MHC-II molecules": Sup Figures 1-7

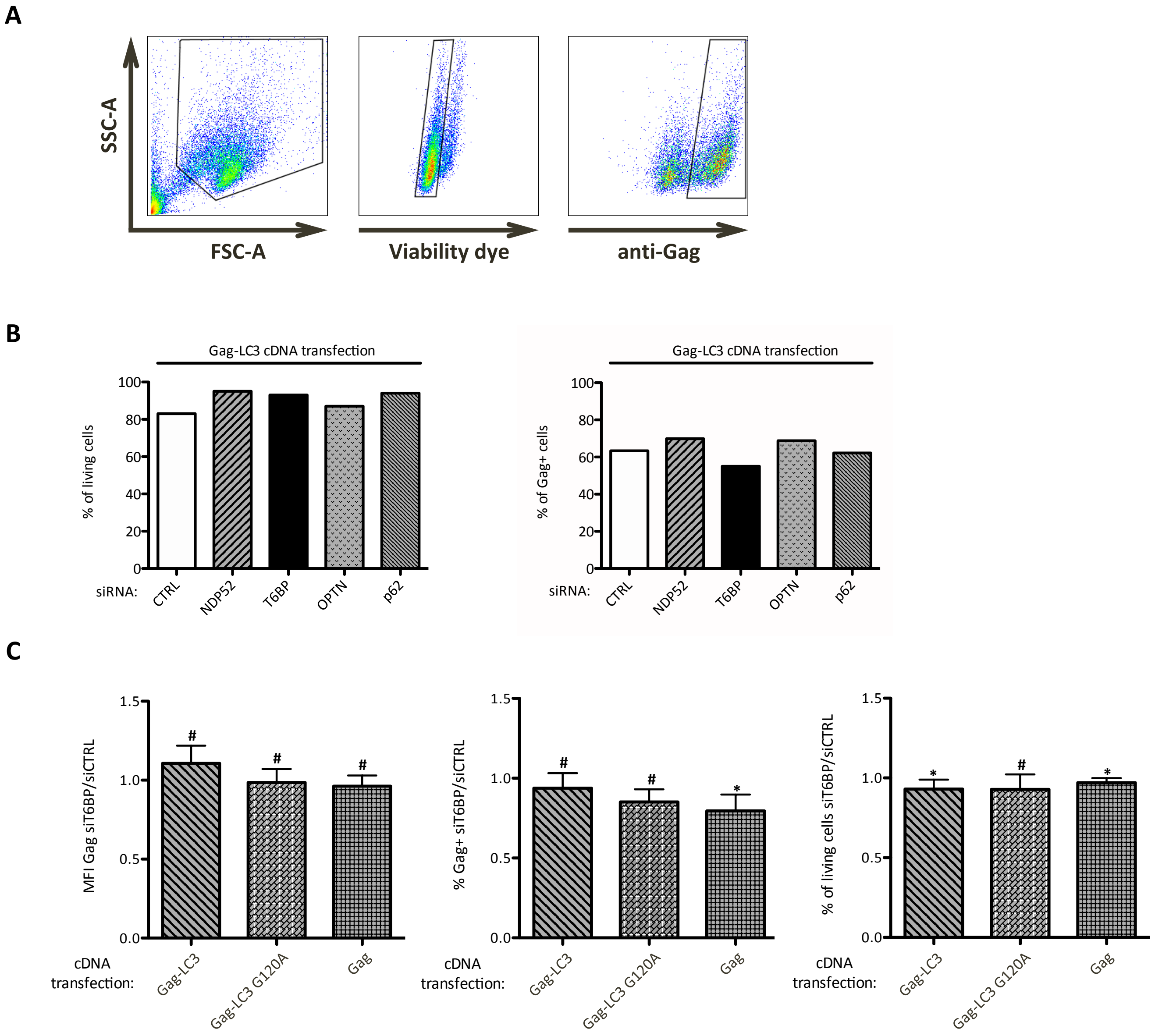


**Figure S1. The transfection of siRNA targeting autophagy receptors influenced neither the cell viability nor the antigen transfection efficiencies. (A) and (B)** Analysis of cell viability and transfection efficiency using flow cytometry. Results of one representative experiment out of three are shown. **(A)** Gating strategy: left panel: FSC-A: forward-scatter; SSC-A: side-scatter; middle panel: staining of living cells using a viability dye; right panel: staining of $Gag^{+}$ cells using anti-Gag antibody intracellular staining. **(B)** Percentage of living and $Gag^{+}$ cells in the different siRNA transfection conditions, left and right panel respectively, analyzed 48h post-treatment, prior co-culture with Gag-specific T cells. **(C)** Results of at least three independent experiments are normalized to control conditions and presented as mean (+/- SD). Left panel, ratio of mean fluorescent intensities (MFI) of Gag stainings in $Gag^{+}$ cells. Middle panel, ratio of the mean percentage of $Gag^{+}$ cells. Right panel, ratio of the mean percentage of living cells. CTRL: control. Mann-Whitney's tests; $*p<0.05$; $\#p>0.05$.

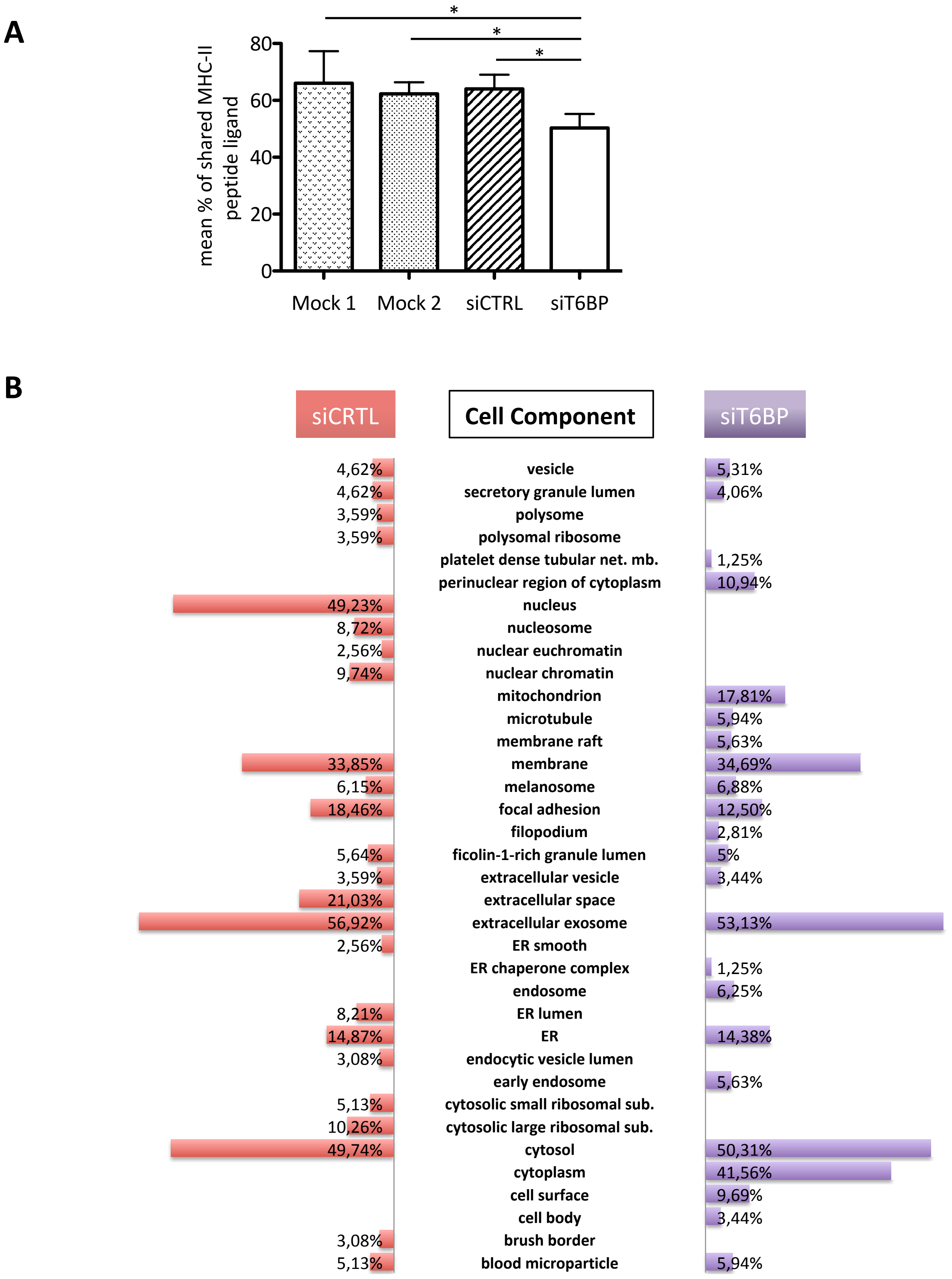


**Figure S2 (related to Figure-3): T6BP silencing affects the repertoire of peptide presented by MHC-II molecules and has a modest influence on the source of MHC-II ligands.** (A) For each sample the % of MHC-II peptide ligand shared with the 3 other experimental conditions was determined and the mean % of shared MHC-II peptide ligand calculated. Comparing the mean % of shared peptides between mock treat cells (Mock1 or Mock2) and the cells transfected with the control (siCTRL) siRNA, no significant differences were observed. In contrast, the mean % of shared MHC-II peptide ligand was significantly different between siT6BP treated cells and Mock1, Mock2 and siCTRL treated cells. The statistical significance was calculated using a Kruskal-Wallis test followed by a Dunn's test (*p<0.05). (B) **Cell component enrichment analysis of peptide sources**. **As in Figure 3,** HeLa-CIITA cells were transfected with siCTRL and siT6BP siRNA, lysed, submitted to MHC-II immunoprecipitation using TÜ39 antibody and the peptide-ligands sequenced using mass-spectrometry (LC-MS/MS). The diversity of protein sources was analyzed according to cell component enrichment using Funrich software. Only canonical pathways statistically enriched (p>0,05) for each conditions (siCTRL and siT6BP) are shown. The p value for pathway enrichment were obtained in right-tailed Fisher's exact tests.

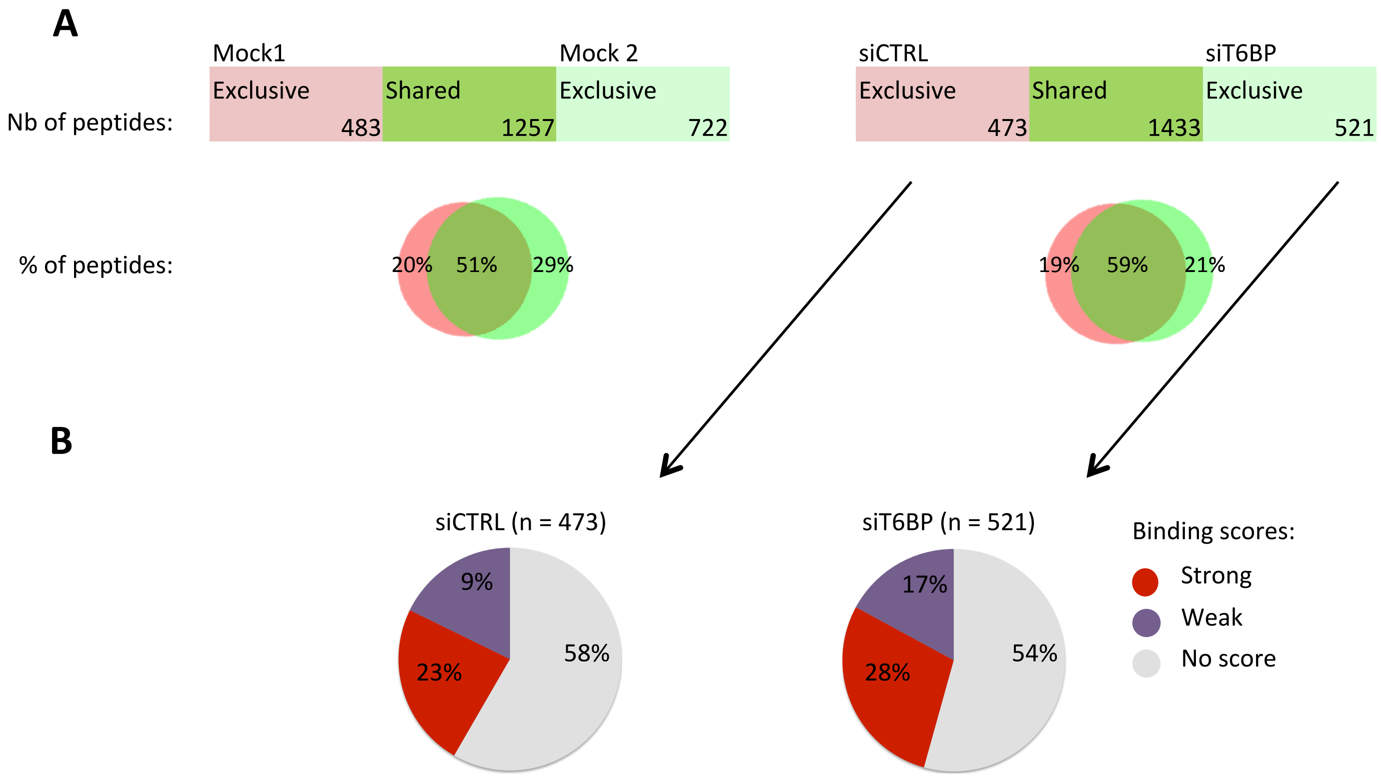


**Figure S3 (related to Figure-3): T6BP silencing does not influence the immunopeptidome of MHC-I molecules. (A) Left panel**, mock-treated HeLa-CIITA cells were split and culture for 48h (giving rise to Mock1 and Mock2), then cells were lysed, MHC-I molecules were immunoprecipitated using W632 antibody and the peptide-ligands sequenced using mass-spectrometry (LC-MS/MS). **Right panel,** HeLa-CIITA cells were transfected with siCTRL and siT6BP siRNA and were treated as in the left panel. The number and the percentage among sequenced peptides (Venn diagrams) of exclusive or shared peptides for each conditions are presented. **(B)** Relative binding affinities, presented as pie charts, of exclusive peptides identified in siCTRL (left) and siT6BP (right) condition (number of peptides are indicated in brackets). NetMHCpan 4.0 algorithm was used to predict the relative affinities to HLA-A*6802 and -B*15093 molecules expressed by HeLa-CIITA cells. The relative affinities to the HLA-C*1203 molecule also expressed by HeLa-CIITA cells were not combined in this figure because many peptides binding to HLA-C*1203 also bind to HLA-A*6802. The results are presented as stated from NetMHCpan 4.0 analysis as Strong (for strong binders), Weak (for weak binders), and No score (for epitopes with which a binding score cannot be determined). Except for the Mock conditions, one representative experiment is shown out of two biological replicates. For each experiment, 5 technical replicates per sample were ran on the LC-MS/MS. Nb : number; % : percentage.

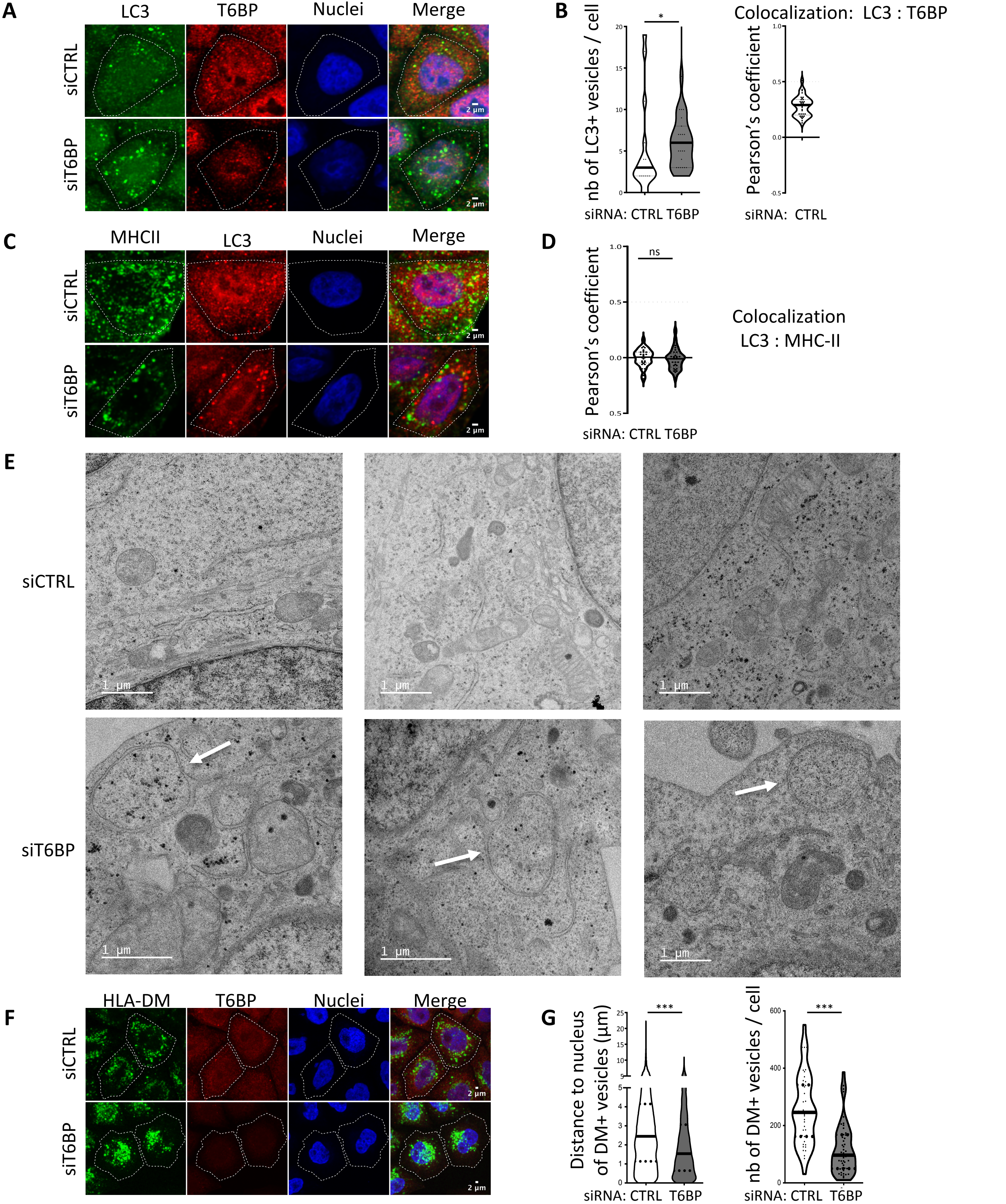


**Figure S4 (related to Figure 4). T6BP silencing leads to autophagosome accumulation. (A)** LC3 and T6BP expressions were assessed using confocal microscopy. HeLa-CIITA cells were transfected with siCTRL or siT6BP. 48h post-treatment, LC3 and T6BP were detected using anti-LC3 and anti-T6BP antibodies, respectively. **B)** Quantitative analysis using in-house ImageJ script displaying the number of LC3$^{+}$ vesicles per cell and colocalization using Pearson's coefficient of T6BP and LC3 staining (right panel). Number of cells = 30. **(C) and (D)** As in A and B with MHC-II molecule and LC3 staining. Number of cells > 40. Scale bars, 2µm. For Pearson's coefficient, the dotted lines (at 0.5) indicate the limit under which no significant co-localization is measured. **(E)** siRNA-treated cells were also analyzed using electron microscopy. Top panels and bottom panels, images from siCTRL- and siT6BP-treated cells, respectively, from 6 representative cells. Two independent experiments were performed and at least 40 cells for each treatment were analyzed. The white arrows indicate the autophagosomes. Scale bars, 1µm. **(F)** As in (A) with HLA-DM and T6BP staining. (**G)** The localization of HLA-DM$^{+}$ vesicles and the number of vesicles/cell were quantified as in Figure-4. At least 10000 vesicles in at least 40 cells were analyzed. Scale bars, 2µm. In graphs representing the number of vesicles/cell, each dot corresponds to a single cell. CTRL: control; nb: number. Whitney's tests; *p<0.05; ***p<0.0001; ns>0.05.

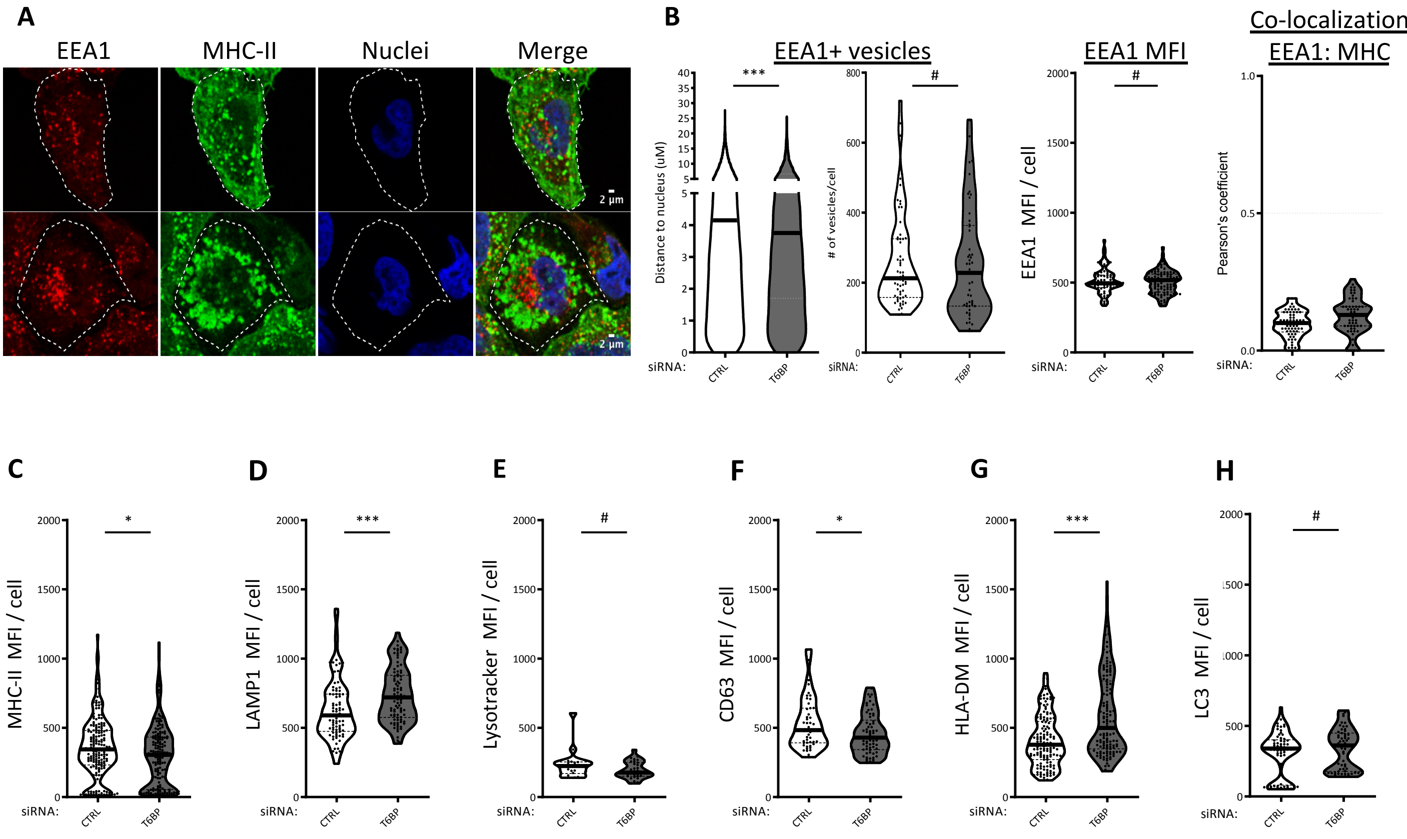


**Figure S5 (related to Figure-4). T6BP silencing slightly influences EEA1+ vesicle cellular localization (A)** EEA1 expression was assessed using confocal microscopy. HeLa-CIITA cells were transfected with siCTRL and siT6BP. 48h post-treatment, EEA1 were detected using specific antibody and a fluorescent secondary antibody. Nuclei were stained using DAPI. **(B)** Quantitative analysis using in-house ImageJ script displaying distance of each EEA1$^{+}$ vesicles to the nucleus, number of EEA1$^{+}$ vesicles per cell and EEA1 total MFI per cell. At least 16 000 vesicles from 90 cells corresponding to 3 independent experiments were analyzed. **(B, Right panel) EEA1 and MHC-II stainings do not colocalize.** Quantification of the potential colocalization between MHC-II$^{+}$ and EEA1$^{+}$ dots using Pearson's coefficient where the dotted lines (at 0.5) indicate the limit under which no significant co-localization is measured. **(C-H) Effect of siT6BP silencing on the expression levels of the various vesicular markers.** As in A, HeLa-CIITA cells were transfected with siCTRL and siT6BP. 48h post-treatment, cells were fixed and labeled and the MFI of the indicated markers were analyzed. **(C)** MHC-II, at least 170 cells corresponding to 5 independent experiments were analyzed. **(D)** LAMP-1, at least 85 cells corresponding to 3 independent experiments were analyzed. **(E)** Lysotracker, at least 30 cells corresponding to 2 independent experiments were analyzed. **(F)** CD63, at least 60 cells corresponding to 2 independent experiments were analyzed. **(G)** HLA-DM, at least 140 cells corresponding to 3 independent experiments were analyzed. **(H)** LC3, at least 60 cells corresponding to 2 independent experiments were analyzed. In graphs representing the number of vesicles or the MFI per cell, each dot displayed corresponds to a single cell. Scale bars, 2µm. CTRL: control. Mann-Whitney's tests; *: $p<0.05$; **: $p<0.002$; ***: $p<0.0003$; #: $p>0.05$.

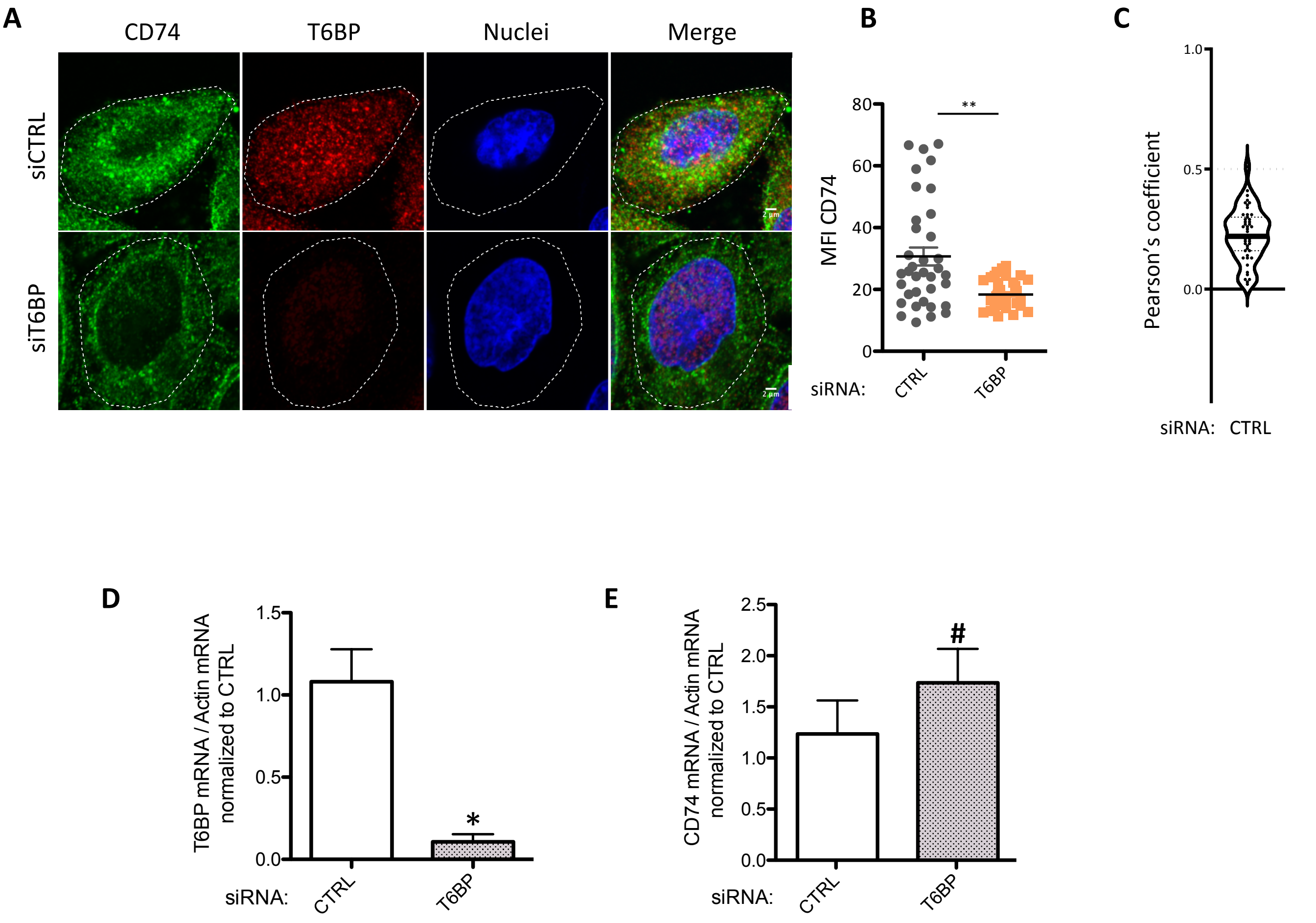


**Figure S6. T6BP silencing affects CD74 expression levels as assess by confocal microscopy but does not affect CD74 mRNA levels. (A)** CD74 expression assessed using confocal microscopy in HeLa-CIITA cells, 48h post treatment with the indicated siRNA. Top panels siCTRL and bottom panels siT6BP. **(B)** Quantitative analysis using ImageJ of CD74 Mean Fluorescent Intensity (MFI). The data are representative of at least 3 independent experiments. Each dot displayed corresponds to a single cell. At least 75 cells were analyzed **(C)** Co-localization of CD74 and T6BP assessed, in the control condition, using Pearson's coefficient. Number of cells = 47. Scale bars, 2µm. CTRL: control. Wilcoxon's tests; **:p<0.002. **(D)** T6BP and **(E)** CD74 mRNA levels were assessed using RT-qPCR. HeLa-CIITA cells were transfected with siCTRL and siT6BP. 48h post treatment, relative T6BP (D) and CD74 (E) mRNA expression levels were analyzed by RT-qPCR using actin as reference gene. Results are presented as a ratio of T6BP (D) and CD74 mRNA (E) levels to actin mRNA level and are representative of four independent experiments. TRL: control. Mann Whitney test: *p<0.05; #p>0.05.

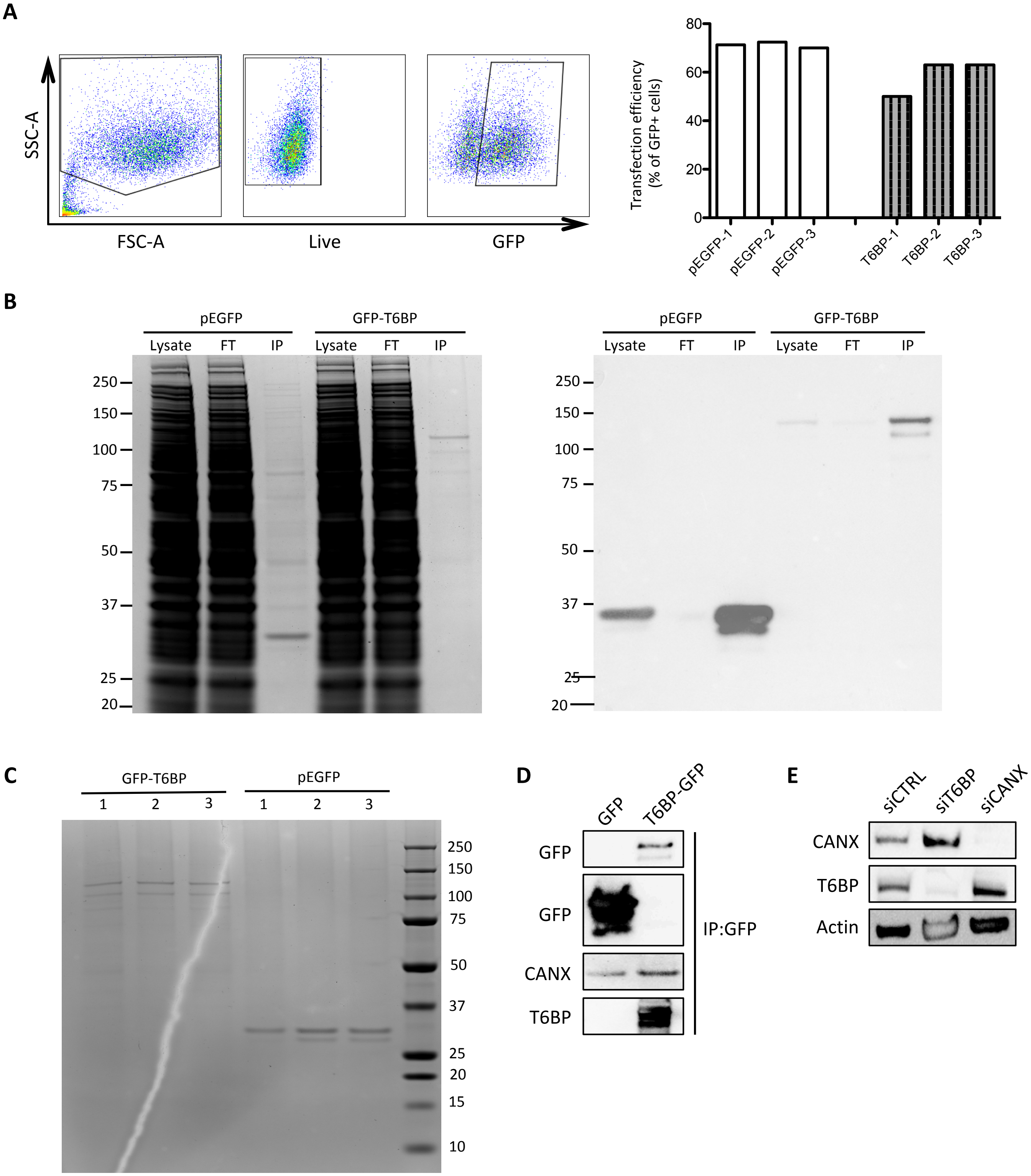


**Figure-S7 (related to Figure-6). Quality controls of T6BP interactome definition. (A)** Left panels, gating strategy for the analysis of GFP⁺ cells after transfection of cDNA encoding wild-type T6BP fused to GFP using flow cytometry. FSC: forward-scatter; SSC: side-scatter. Right panels, side to side comparison of the percentage of GFP⁺ HeLa-CIITA cells transfected with plasmid encoding GFP-T6BP or GFP-only (pEGFP) in the three biological replicates. **(B)** Coomassie blue staining (left panel) and Western Blot analysis (right panel) of the indicated fraction of protein samples from GFP-T6BP and GFP expressing cells. Results from one of the three biological replicates are shown. Briefly, cells were lysed and submitted to IP using anti-GFP camel antibodies (GFP-Trap from Chromotek). Right panel, GFP and GFP-T6BP were revealed using anti-GFP antibody. FT = Flow through. IP = Immunoprecipitation. **(C)** Coomassie blue staining of immunoprecipitated proteins (1/10 of the sample volume) used for LC-MS/MS analysis (9/10 of the sample volume) of the three biological replicates for both GFP-T6BP and GFP. **(D)** GFP nanobody immunoprecipitates from HeLa-CIITA cells transfected with GFP and T6BP-GFP. 48h post-transfection, samples were analyzed by Western blot with the indicated antibodies. **(E)** Silencing of T6BP expression does not influence calnexin (CANX) expression levels. HeLa-CIITA cells transfected with the indicated siRNAs and samples analyzed, 48h post transfection, by Western blot with the indicated antibodies.
